# Multidimensional single-cell modeling of cellular signaling

**DOI:** 10.1101/2020.11.15.383711

**Authors:** James D. Wade, Xiao-Kang Lun, Bernd Bodenmiller, Eberhard O. Voit

## Abstract

Cell-to-cell differences in signaling components can lead to qualitatively different responses to stimuli. Understanding this heterogeneity in signaling response is limited by the inability of time-lapse methods to measure multiple pathway components simultaneously in situ. Here, we present *Distribution-Independent Single-Cell ODE modeling* (DISCO), a computational method for inference of continuous single-cell signaling dynamics from multiplexed snapshot data. We used DISCO to analyze signaling in the MAPK/ERK pathway of HEK293T cells stimulated with the growth factor EGF. Our model recapitulates known features of the ERK signaling response and enables the detection of hidden cell-to-cell variation in seemingly homogeneous samples. Further, DISCO analysis suggested that the MAPK/ERK pathway transmits signal duration rather than amplitude, and that cell-to-cell variation in MAPK/ERK signaling response depends primarily on initial cell states. Finally, we applied an extended version of DISCO to explain changes in signaling kinetics due to overexpression of a disease-relevant protein. Overall, DISCO enables a deeper understanding of how single-cell variation affects cellular responses in complex signaling systems.

## Introduction

Cell signaling pathways are complex biochemical systems at the core of cellular information processing. The dynamics of these signaling systems in response to extracellular cues is critical for proper cell functioning (Dolmetsch et al., 1997; Kholodenko, 2006; Selimkhanov et al., 2014). Signals are transduced by modulating enzymatic activities and local concentrations of signaling mediators such as protein kinases. Cell-to-cell variation in expression of signaling components, even within a clonal cell population, can lead to different responses and functional outcomes, such as proliferation instead of apoptosis (Spencer et al., 2009). In cancer and other diseases, genetic alterations change the expression or function of signaling components, increasing cellular variation and creating cells with aberrant signaling responses compared to healthy cells (Altschuler and Wu, 2010). In addition, microenvironmental differences can further diversify cancer cell cell states and, consequently, drug responses (Meacham and Morrison, 2013; Burrell and Swanton, 2014). In such cases, one subgroup of cells may strongly respond to a particular drug treatment while another is insensitive (Burrell and Swanton, 2014).

To characterize cell-to-cell variation in signaling dynamics, the changes in abundances or, ideally, concentrations of signaling molecules within individual cells must be observed over time. This is experimentally possible through live-cell imaging methods such as Förster resonance energy transfer, kinase translocation reporters, and activity-dependent dyes that measure the activation state or activity of signaling molecules (Ryu et al., 2015; Regot et al., 2014; Dolmetsch et al., 1997). However, cell signaling networks consist of numerous interacting components, whereas even multiplexed livecell methods are currently limited to observations of only a few pathway components at a time (Bunt and Wouters, 2017; Regot et al., 2014). As an alternative, multiplexed single-cell methods, such as fluorescent and mass cytometry, provide a large dynamic range and can simultaneously measure up to 50 signaling network components (Lin et al., 2015; Giesen et al., 2014; Bodenmiller et al., 2012; Lun et al., 2017; Gut et al., 2018). As these methods require that cells be fixed at a point in time, they provide only snapshots of single-cell states. Thus, although these highly multiplexed methods can be used to more comprehensively characterize signaling networks, they cannot be used to follow the signaling dynamics of individual cells over time, so that the critical link between initial cell state and response to stimulus is lost.

Computational methods have been used at the population level to overcome the trade-off between continuous and snapshot measurements. In such approaches, a mathematical model of the underlying system is used to capture the range of potential system behaviors. The biochemical reactions of cell signaling are commonly modeled using a system of deterministic ordinary differential equations (ODEs) (Wang et al., 2015; Filippi et al., 2016). Forward-simulation of the ODE model generates continuous trajectories of state variables, such as the average phosphorylation states of signaling proteins. System parameters that define specific system trajectories are inferred by matching simulations to snapshot measurements.

Given the well-established importance of cell-to-cell variations in signaling responses (Spencer et al., 2009; Altschuler and Wu, 2010; Meacham and Morrison, 2013; Burrell and Swanton, 2014), modeling approaches that specifically leverage single-cell measurements are needed (Bronstein et al., 2015). Previous methods to account for cellular variation in signaling dynamics have used parametric distributions (i.e., distributions defined by a fixed set of parameters such as the mean and standard deviation) to represent single-cell snapshot data instead of simulating individual cell states (Hasenauer et al., 2011; Hasenauer et al., 2014; Filippi et al., 2016; Loos et al., 2018). Use of parametric distributions, however, complicates model inference. The form of the distribution, such as Gaussian or gamma, and associated parameter values, must be chosen to approximate the experimental samples prior to kinetic modeling. More importantly, a single parametric distribution is unlikely to be a good approximation of samples that contain multiple populations (Fig. 1), and additional assumptions and parameters are needed to represent the number and relative mixing components of sub-populations. As the number of distribution parameters rapidly increases with the number of system state variables (e.g., signaling network components) and/or as samples contain increasingly complex mixtures of subpopulations, parametric formulations rapidly become analytically intractable. These challenges have been addressed by ignoring covariance of variables (Hasenauer et al., 2014; Filippi et al., 2016) or by considering distributions with no more than two measurement dimensions (Loos et al., 2018), thus reducing the number of parameters but resulting in a loss of valuable information unique to multivariate single-cell data (Sachs et al., 2005). Importantly, if the assumptions used to represent samples are not accurate, the inferences are likely to be incorrect. In recent work, a maximum entropy-based approach to infer cell-to-cell heterogeneity in signaling network parameter was reported, and applied to study response heterogeneity in EGFR/Akt signaling (Dixit et al., 2020). This approach however is most appropriate for 1-dimensional data and there remains a need for methods to infer signaling responses from multidimensional data, where multiple markers are monitored simultaneously.

**Figure 1:**
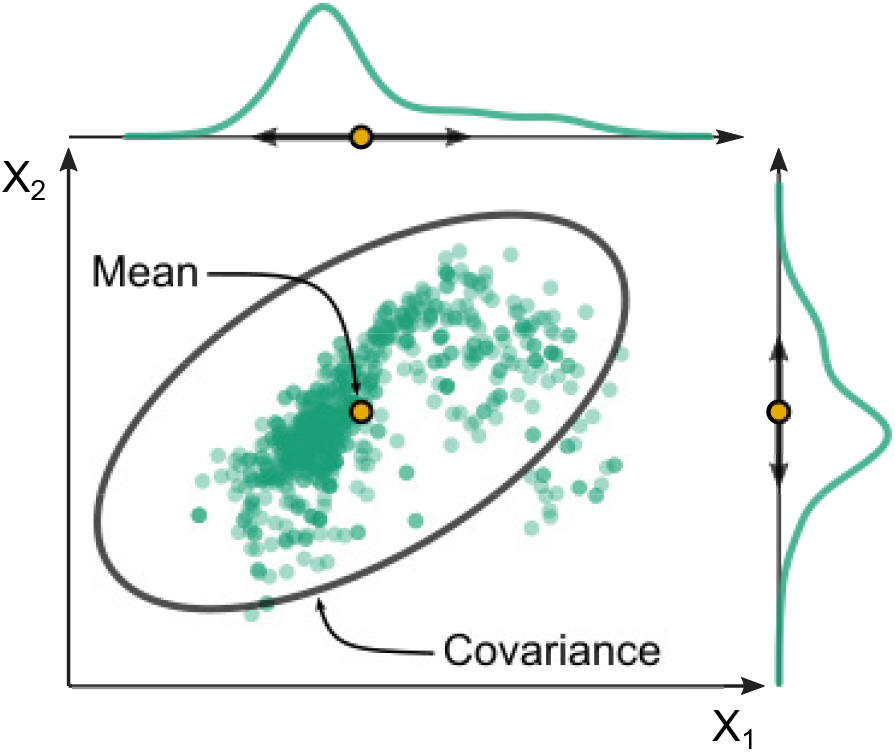
Disadvantage of representing single-cell data by parametric distributions. An example single-cell measurement (green dots). Approximating the cell population by the mean (yellow circle) and covariance (black ellipse) of a log-normal distribution fails to capture the complex underlying biological structure.

Here we describe a novel computational approach for inference of multiplexed single-cell signaling trajectories from snapshot data. Our methodology uses an ODE system to simulate the trajectories of many individual cells and applies a distribution-free statistical test to compare experimental and simulated snapshots and quantify model fitness. This approach overcomes the challenge of determining the number, distribution shape and associated parameters of cell populations by using the single cell observations as a native representation of the biological variation in the sample.

We used our approach to analyze single-cell variation within the well-studied MAPK/ERK signaling cascade in HEK293T cells stimulated with the growth factor EGF and monitored by multiplexed mass cytometry of 12-16 cell-state variables. Our approach reproduces known features of ERK signaling and enables analyses of features that cannot be measured directly, such as the relationship between signal duration and amplitude throughout the pathway. Our model enables formulation of several hypotheses for future experimental validation: it suggests that single-cell variation in immediate signaling responses can be described deterministically and that the pathway serially transmits signal duration at the single-cell level. It also helps us identify activated sub-populations within seemingly homogenous samples, illustrating that cell-to-cell differences not observed experimentally can still be inferred. Finally, as progression of many cancers is related to protein overexpression (Santarius et al., 2010), we apply our approach to samples overexpressing disease-related proteins and observe that protein overexpression alters the modeled reaction kinetics in a nonlinear fashion. Overall, we present a method that enables inference of continuous dynamics from discrete snapshot data in heterogeneous systems and quantifies variation in cellular response at the single-cell level.

## Results

### DISCO infers dynamic signaling responses from multiplexed single-cell snapshots

Determining sources of cell-to-cell variation in signaling responses requires two components: singlecell signaling trajectories and cell-state variables that determine signaling responses. Current experimental methods cannot capture both simultaneously: methods that allow measurement of singlecell trajectories generally lack the dimensionality to observe all the variables driving differential response (Fig. 2a), whereas higher-dimensional snapshot measurements lack the time-dependent single-cell dynamics needed to quantify response (Fig. 2b).

**Figure 2:**
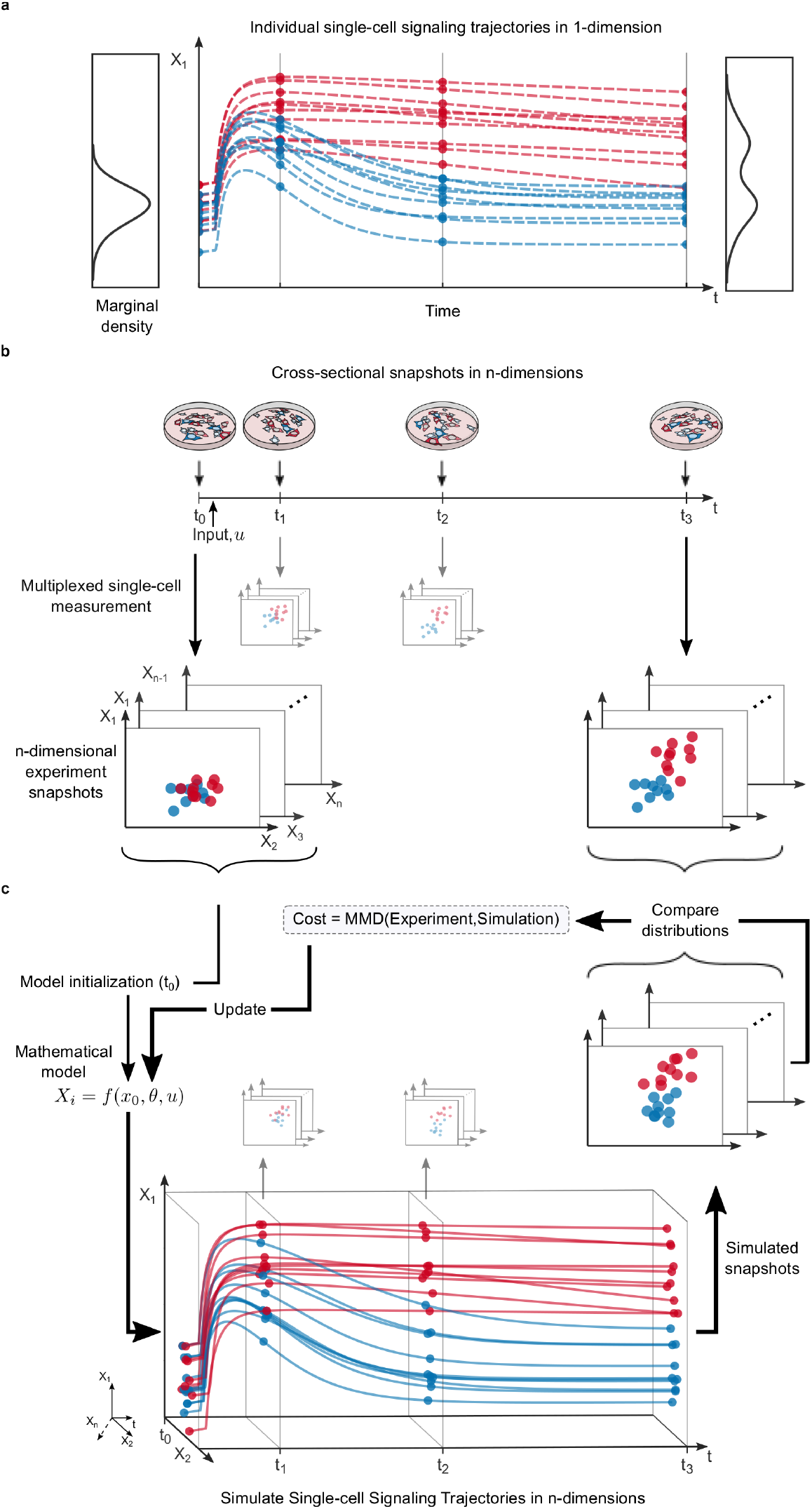
Motivation and workflow of DISCO. (**a**) Motivation. One-dimensional time-lapse measurements (dots) of variable a (X_1_) do not characterize the source of a bimodal signaling response. Red and blue trajectories represent distinct subpopulations. (**b**) Experimental workflow. Multiplexed experimental snapshots are taken of many single cells responding to a perturbation over time. Snapshots capture multiple markers (X_1_,…, X_*n*_), but cannot measure response trajectories. For visualization purposes, scatter plots represent illustrative two-dimensional projections of n-dimensional distribution snapshots. (**c**) Computational workflow. Steady-state measurements at to are used to initialize a set of ODE model instances, one for each cell. In this case, *X_i_, i* = {1, 2} is shown for illustration. Differences between simulated and measured snapshots of n-dimensional cell-state distributions are compared and used to optimize model parameters. This is done for all experimental time points, t_1_,…, t_*n*_, shown for simplicity with smaller insets.

To characterize continuous multiplexed single-cell signaling trajectories and study the origins of cell-to-cell differences in signaling response, we developed a novel framework combining experimental and computational methods, called *Distribution-Independent Single-Cell ODE modeling* or DISCO (Fig. 2b-c). This approach yields continuous multidimensional single-cell trajectories bridging the gap between continuous and snapshot measurements.

DISCO uses an ODE system to describe the trajectories of individual cells, so that each cell has its own parameter set and model-based trajectory. These individual trajectories are then combined to represent the trajectory of a an empirical distribution that describes the sample, which is then compared to experimental data using non-parametric statistical methods. To quantify the distance between experimental and simulated snapshots and thus fit the model, we used maximum mean discrepancy (MMD) (Gretton et al., 2012), a kernel-based, distribution-free, two-sample test for multivariate distributions. The use of distribution-free statistics for model fitting has major advantages over parametric approaches for modeling of cellular variation in signaling. First, it decouples the number of free model parameters from the number of cells or subpopulations, which makes any single-cell modeling problem similar in parameter number to a corresponding classical population-average model. Second, it can successfully discriminate between differences in distributions otherwise missed by parametric statistical methods, since it does not make assumptions about the shapes or numbers of cell populations. Finally, DISCO is readily integrated into classical modeling frameworks: single-cell models are instantiated using single-cell measurements, and simulation of many cells fits within ensemble modeling frameworks. We also found, through a test case used by parametric approaches (Hasenauer et al., 2014; Loos et al., 2018), that our method successfully extracts the parameters of latent cell populations (For a full analysis, see the Supplementary Notes).

#### Single-cell variation in MAPK/ERK signaling can be described deterministically

We chose to test the DISCO approach on the well-studied MAPK/ERK signaling cascade because known features of this pathway can be used to test the validity of the model inferences. Additionally, expression and/or function of ERK pathway components is deregulated in many cancers, the pathway is a common target of cancer drugs (Samatar and Poulikakos, 2014; Caunt et al., 2015), and cell-to-cell variation has limited the success of these targeted therapies (Wellbrock and Arozarena, 2016; Caunt et al., 2015).

We used mass cytometry to measure, in triplicate, multi-dimensional single-cell snapshots of cell states. We measured total and phosphorylated (active) MEK, ERK, and p90RSK, as well as cellcycle, death, and cell-size markers at six time points in HEK293T cells (on average >10,000 cells per sample) stimulated with EGF over a 1-hour time course. Then, we used DISCO to construct a single-cell model of the pathway (Fig. 3a). While the true pathway involves many more reactions, we condensed it to a core pathway that includes active RAF and both the active and inactive states of MEK and ERK. The model also includes active and inactive p90RSK, which is a downstream target of ERK, and GAPDH, which is modeled as a constant for each cell (Fig. 3a). For model fitting, we used measurements of total and active MEK, ERK, and p90RSK, but not GAPDH, from cells subsampled across the experimental replicates at six time points.

**Figure 3:**
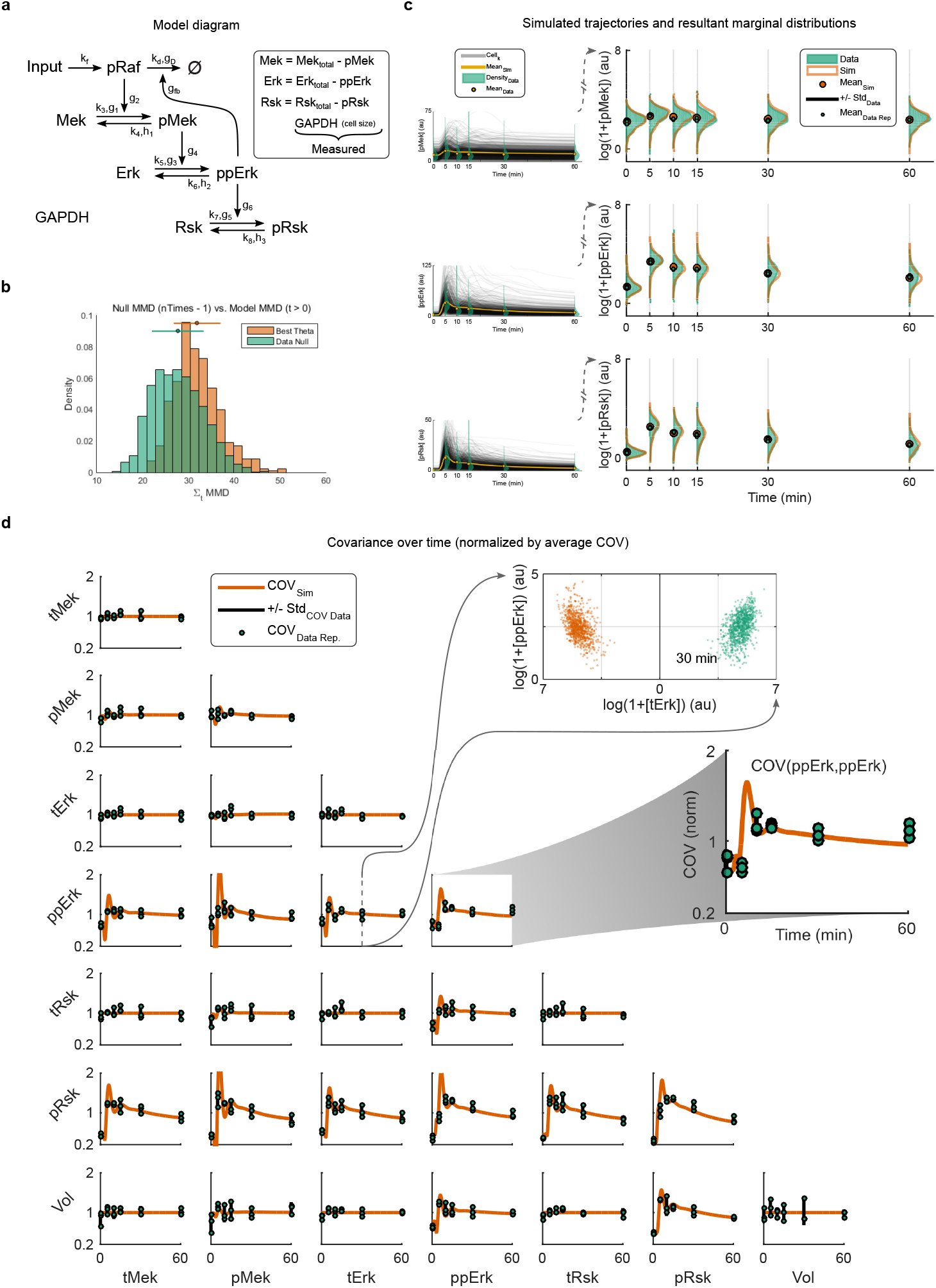
Single-cell model of the RAF-MEK-ERK-p90RSK pathway. (**a**) Diagram of the pathway model with kinetic parameters, which are inferred from measurements of total and active forms of MEK, ERK and p90RSK. GADPH is represented as a constant, not used in fitting and therefore measured for validation purposes. (**b**) Comparison of experiment-based (null) and model-based MMD distributions excluding *t* = 0 min. (**c**) Insets on left: Simulated single-cell trajectories (black lines) compared to snapshot measurements (green histograms). Yellow circles and line are the mean of the data and simulations, respectively; note linear y-axis. Right: Marginal (one-dimensional) distributions of measurement data (green) and single-cell model simulations (orange); note log y-axes. (**d**) Covariance between model components over time for measurements (green circles; by replicate) and model simulations (orange line). Vertical black lines represent the covariance averaged across replicates +/− the standard deviation in covariance across replicates. (Top inset) Example of a scatter plot comparing single-cell data (green) and single-cell simulations (orange) at a given time point. The x-axis is mirrored about 0, so that a green-orange mirror image would indicate a perfect model fit. (Lower inset) Zoomed example of variance of ppErk over time. In each plot, covariance has been normalized by the average covariance over time.

To quantitatively assess model fit, we used bootstrapping to compare a null MMD distribution based on experiments to a distribution of the MMD statistic generated by simulations with different cell subsamples. Model parameters remained fixed across samples. Comparison of the null (mean = 27.4; std = 5.5) and model (mean = 31.9; std = 5.0) MMD distributions showed that the overall model fit was within range of experimental variation (Fig. 3b). This was also the case at the level of individual times points (Supplementary Fig. S6). Comparison of model simulations to data showed that the model successfully captured the changes in population statistics, such as the mean and covariance, even though it was not explicitly fit to these values. In addition, the model captures the shape of the marginal distributions and higher dimensional distribution structures in the data (Fig. 3c-d). Thus, our modeling approach is capable of capturing cell-to-cell signaling variation in a population of cells, and this variation can be described as a deterministic function of initial cell state.

**Figure 4:**
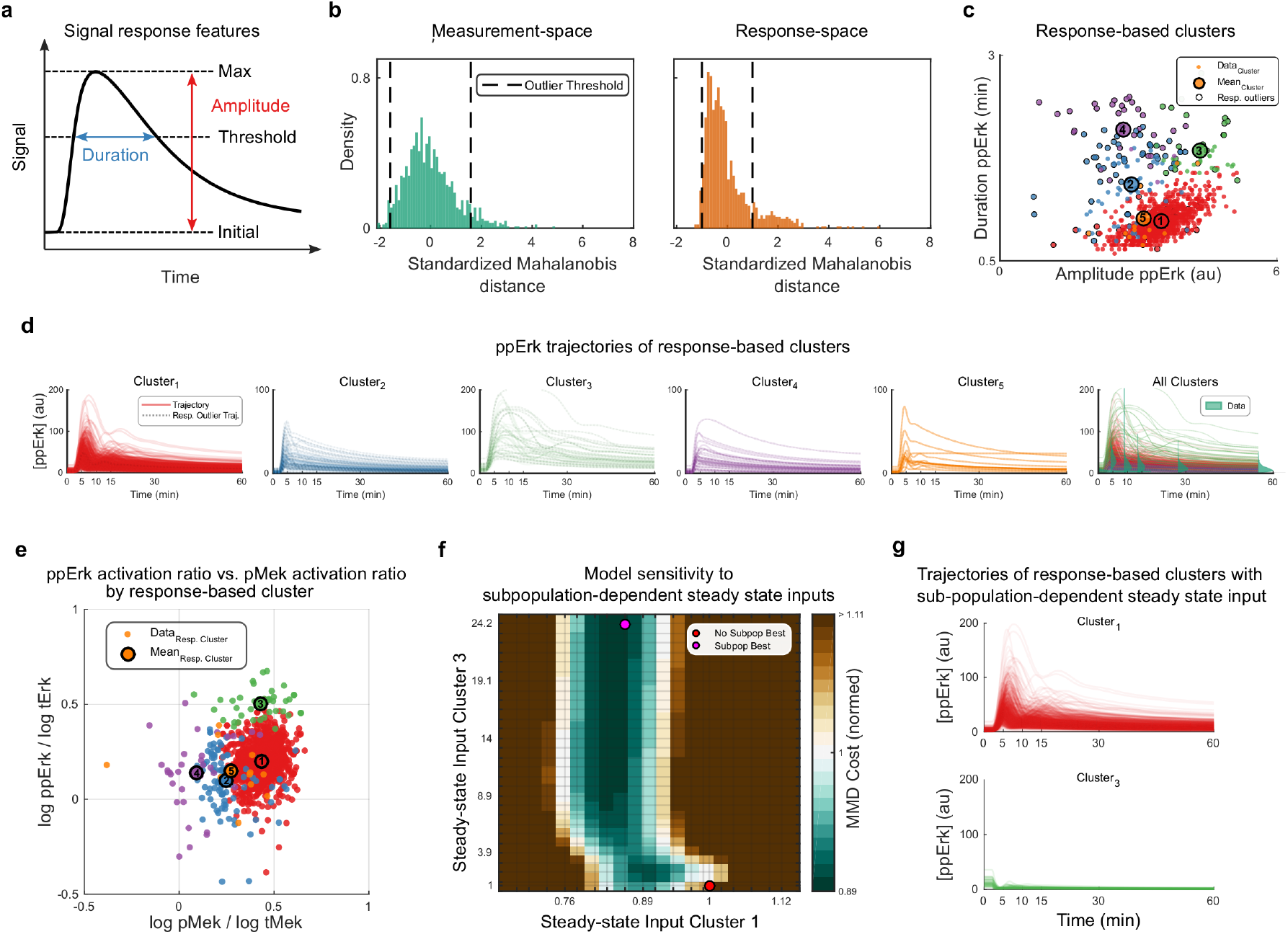
Response-based subpopulations. (**a**) Response features calculated for each inferred individual cell trajectory. (**b**) Distribution of Mahalanobis distances across all cells in measurement space (left) and response space (right). (**c**) Projection of the response-based clusters onto the two dimensions of ppErk amplitude and duration. Cells are colored by cluster ID. Large numbered circles are cluster centers. Response-based outliers are circled in black. (**d**) Trajectories of ppErk for all cells in each response-based cluster. Response-based outliers are overlaid with dashed lines. Note cluster-dependent y-scale. (**e**) Response-based clusters projected in the measurement space. Large numbered circles are cluster centers. (**f**) Model fitness as a function of the steady-state input for Cluster 1 and Cluster 3. Cluster-dependent inputs improve model fitness (magenta circle) compared to population-wide input that assumed a homogenous population with input arbitrarily set to 1 (red circle). (**g**) ppErk trajectories for cells in Response-Clusters 1 and 3 associated with the steady-state input at the magenta circle in (f).

#### Single-cell trajectories correctly predict an additional cell-state dimension

To validate the model, we predicted the multi-dimensional relationship between simulated signaling proteins and a measured cell-state variable not used in model fitting, GAPDH abundance, which is a surrogate for cell volume (Rapsomaniki et al., 2018) (Supplementary Fig. S13). Although cell volume has predictive value for signaling state and could therefore be used to improve model fit, we did not include this information in the model calibration. GAPDH does not change during the 60 minute time-period of the experiment (Supplementary Fig. S13) and should not directly affect the signaling reactions. It can therefore be represented in the model as a cell-specific constant for each cell during simulation (Fig. 3a). The signaling trajectories inferred by the single-cell model predicted the experimentally observed GAPDH levels across all time points (3d). We note that this result is not a guaranteed consequence of model construction as, although GAPDH correlates with variables that were used in parameter fitting, correlation is not a transitive property in multivariate spaces (Castro Sotos et al., 2009). Thus, the ability of the single-cell model to correctly predict the relationships between signaling variables and a measured cell-state variable supports the validity of the inferred single-cell trajectories.

### Multivariate analysis of variation in signaling response of the MAPK/ERK pathway

Variations in ERK signaling have been suggested to result from extrinsic (unmeasured) variation upstream of MEK (Filippi et al., 2016), but the source of variation in signaling remains an open question due to the lack of continuous multiplex measurements. We used the multiplexed singlecell trajectories provided by our model to identify potential sources of variation in ERK pathway signaling. As key metrics of the signaling response (Shaul and Seger, 2007), we calculated peak amplitude and duration (Fig. 4a) of each active pathway component, namely phosphorylated MEK (in the model: pMek), phosphorylated ERK (ppErk), and phosphorylated p90RSK (pRsk). We then analyzed how variation in the measured initial cell-state variables relates to variation in the six resulting response features of the pathway for each cell.

#### Analysis of single-cell response features identifies a hidden cell population

Even seemingly homogenous cell populations can harbor hidden functional heterogeneity, and modeling single-cell trajectories may enable the identification of such latent subpopulations. To identify cells that may belong to hidden populations, we used the distribution of single cell Mahalanobis distances (Mahalanobis, 1936) in a given feature space to define outlier cells. We found that the response space in our model, defined by the amplitude and duration of all signaling variables, clearly contained outlier populations of cells, whereas the measurement space of cell-states at steady state did not (Fig. 4b). To better understand these response-based outliers, we used a regularized Gaussian mixture model to cluster the cells in either the measurement or response space (Supplementary Fig. S7, Fig. 4c, respectively). In the response space, cells were grouped into five clusters with qualitatively observable differences in signaling trajectories (Fig. 4d). Interestingly, Cluster 3 contained the approximately 3% of outlier cell trajectories that were above the range of snapshot measurements at the 15- and 30-minute time points (Fig. 4d). Further analysis revealed that an especially high ratio of initial ppERK to total ERK, but not cell volume, was predictive of Cluster 3 (Fig. 4e, Supplementary Fig. S7) and suggested a potentially more active or pre-activated cell state. These cells were not identified by clustering in the measurement space (Supplementary Fig. S7 and Supplementary Fig. S8), which was likely due to the multivariate and nonlinear nature of the response features calculated from initial cell states.

We used our model to test the idea that variation in the steady-state input, such as receptor level or activation state, could be driving differences between response clusters. Specifically, we allowed the steady-state input, which had been fixed, to vary across cells according to response cluster. Optimization of these five new subpopulation-dependent parameters (one for each of five response clusters with all other model parameters fixed) improved model fitness by approximately 10%, a rather large qualitative increase (Fig. 4f, Supplementary Fig. S9). The primary qualitative change in cluster behavior was in Cluster 3, which went from highly responsive to non-responsive to EGF, supporting the hypothesis that this cluster represents a pre-activated subpopulation of cells that are relatively insensitive to the stimulus (Fig. 4g). These results illustrate how DISCO can be used to identify hidden subpopulations in cells samples that appear homogeneous when judged by measurements alone.

#### MAPK/ERK signaling cascade preferentially transmits signal duration rather than absolute amplitude

We used our single-cell model, including the subpopulation-dependent steady-state input, to analyze cell-to-cell variation in a model-wide manner (Methods). We found that the variation in amplitude of each signaling component is better explained by its initial state than by the amplitude of the upstream activator. Specifically, the initial steady-state levels of pMek, ppErk, and pRsk explained 75%, 48%, and 42% of the variation in amplitude of these markers, respectively. In contrast, pMek amplitude explained 38% of the variation in the amplitude of ppErk, which in turn explained 27% of the variation in pRsk amplitude (Fig. 5, Supplementary Data). In other words, each variable preferentially exhibits a fold-change (amplitude/initial) response rather than a response dependent on the absolute amplitude of its respective activator. This is also illustrated by the reduction in the coefficient of variation (CV), a measure of the normalized distribution width where smaller values represent a relatively more narrow distribution, of the fold-change in amplitude (amplitude/initial state) (CV_*FoldAmp.pMek*_ = 0.28, CV_*FoldAmp.ppErk*_ = 0.26, CV_*FoldAmp.pRsk*_ = 0.32) compared to the CV of the amplitude (CV_*Amp.pMek*_ = 0.69, CV_*Amp.ppErk*_ = 0.65, CV_*Amp.pRsk*_ = 0.67,5b). Notably, the relative reduction in CV between the absolute and fold-change ERK amplitudes agrees with live-cell imaging of the nuclear to cytoplasmic ratio of fluorescently tagged ERK (Cohen-Saidon et al., 2009), which offers additional support of the model-inferred response features.

**Figure 5:**
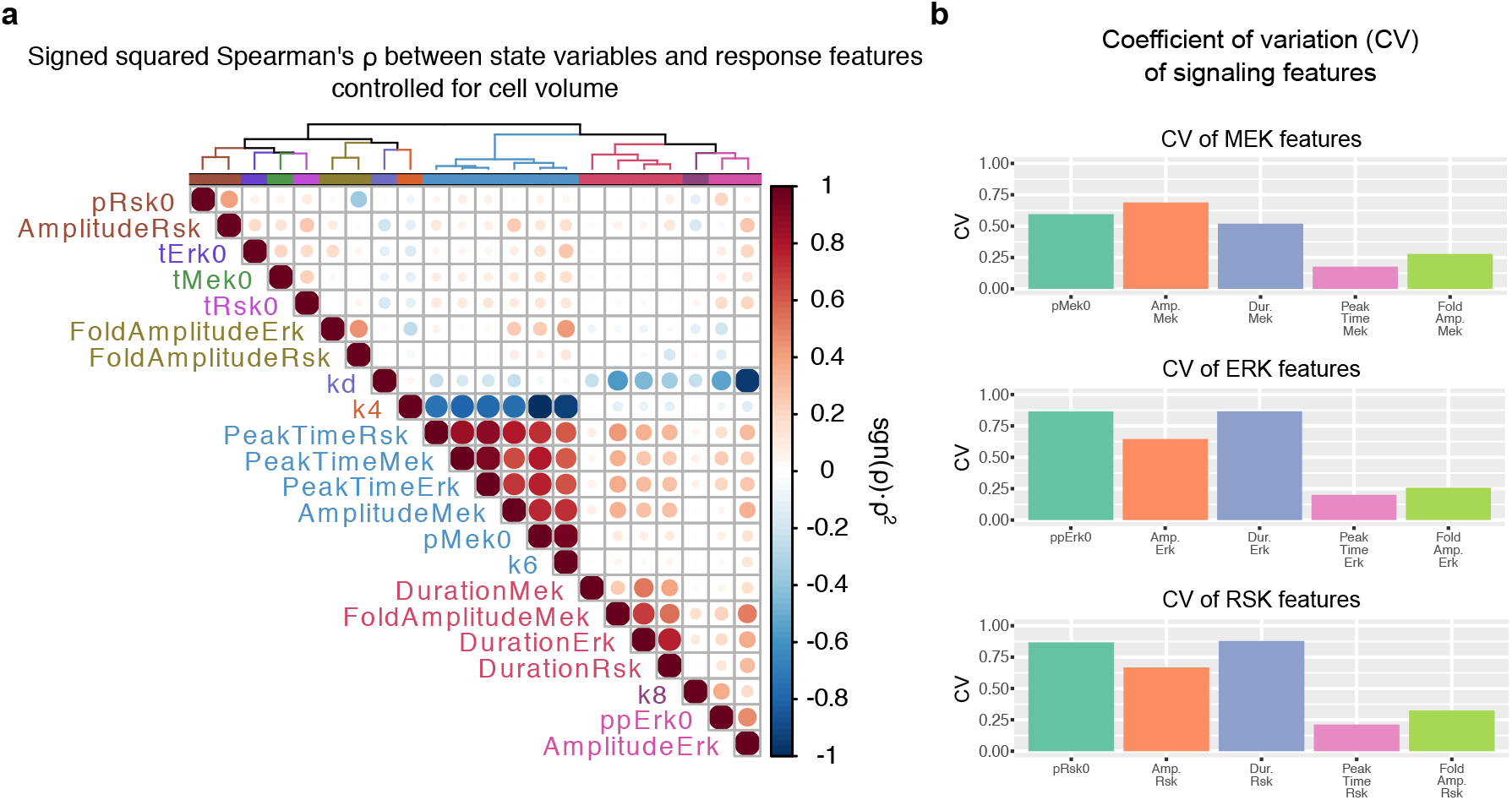
Analysis of multivariate measurement and response features. (**a**) Clustered heatmap of signed squared Spearman’s *ρ* between measured steady-state variables and response features after regressing out cell volume effects. Labels are colored according to dendrogram cut shown. Circle diameter and color illustrate sgn(*ρ*) · *ρ*^2^ as in the colorbar. (**b**) Coefficient of variation calculated for both measurement-and response-based features at each level in the signaling pathway.

When analyzing signal duration, we found high explanatory power moving serially down the pathway. The duration of pMek explained 53% of the variation in the duration of ppErk, which explained 77% of the variation in the duration of pRsk signaling (Fig. 5a). The variation in signal duration across the population, however, (CV_*Dur.pMek*_ = 0.52, CV_*Dur.ppErk*_ = 0.87, CV_*Dur.pRsk*_ = 0.88) was similar to variation in initial cell states (CV_*pMek0*_ = 0.60, CV_*ppErk0*_ = 0.87, CV_*pRsk0*_ = 0.87, 5). Thus, signaling duration is tightly coupled within the pathway in individual cells, and its variation scales with variation in cell state. The combination of population-wide fold-change with single-cell-specific duration suggests that the MAPK/ERK pathway transmits signal duration rather than absolute amplitude.

### Kinetic effects of ERK overexpression on MAPK/ERK signaling

Cancers are caused by genomic alterations such as mutations and copy number alterations (Blume-Jensen and Hunter, 2001) (Abu-Remaileh et al., 2015; Jeong et al., 2014) that greatly increase the range of protein expression in a cell population and thereby alter signaling behavior (Lun et al., 2017). Overexpression of a protein involved in a signaling network should not change the structure of that network, which allows us to hypothesize that overexpression-related signaling changes should be a function of expression differences alone. To test this hypothesis, we used an experimental overexpression system and adapted DISCO to model a system with a large range of expression of network components.

The single-cell model we used up to this point was constructed using non-saturable power functions that are known to represent reaction kinetics in an effective, but minimalistic manner (Savageau, 1976; Voit, 2013). Power functions are local approximations of the “true” reaction speed and well suited for capturing normal operating ranges of variables. For instance, the protein expression data analyzed above were modeled well by the power-law representation. These functions may, however, become too inaccurate for representing very large deviations of expression. For instance, they do not saturate, which is expected to occur in cells under stress, and a sufficient increase in expression will lead to a vastly overestimated reaction kinetics in the model (Fig. 6a). We therefore expanded our modeling approach to infer overexpression-related kinetic changes.

**Figure 6:**
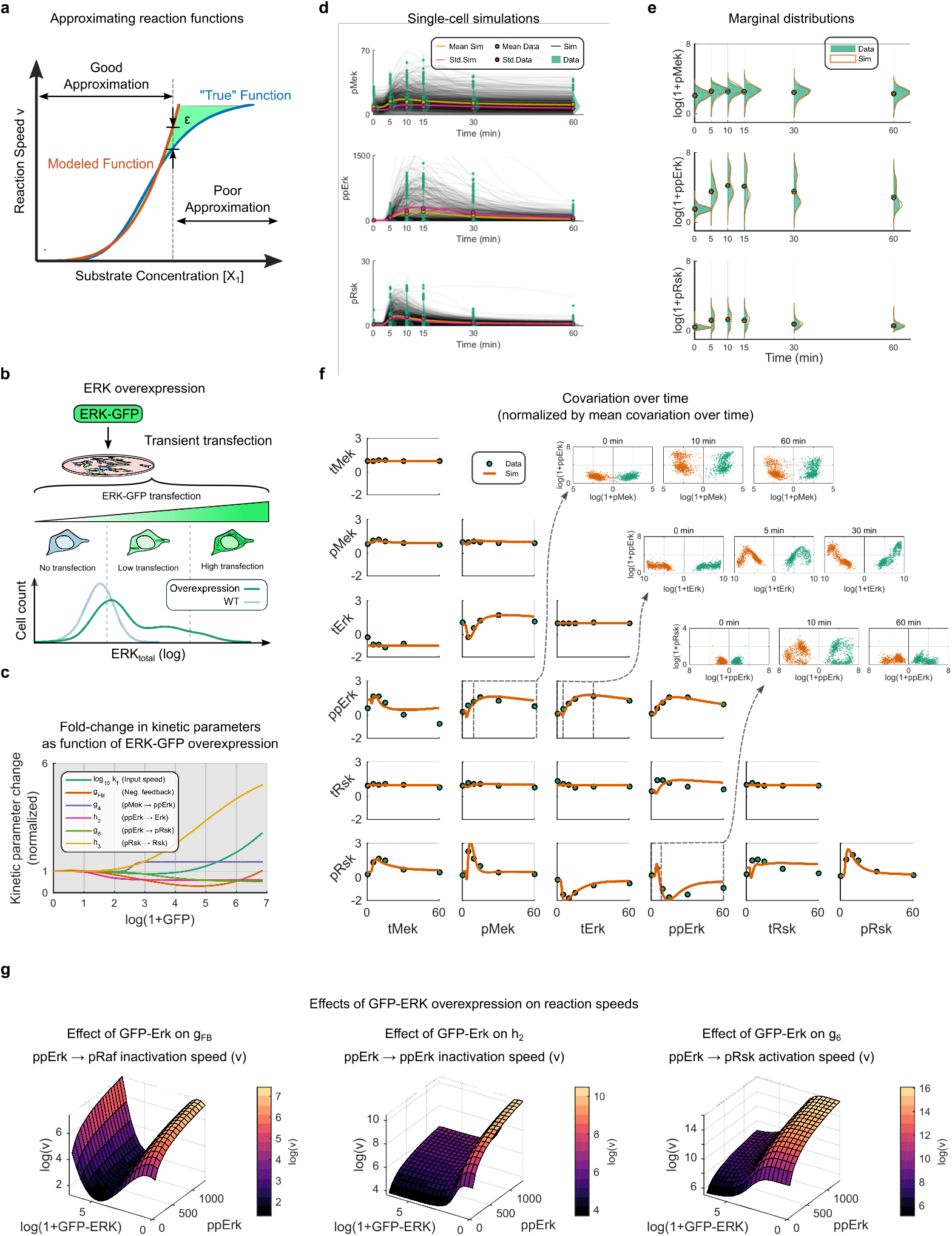
Inferring the effects of overexpression on reaction kinetics. (**a**) Substantially increasing the range of expression can reveal changes in kinetic functions that are not otherwise observable. (**b**) Transient transfection induces a continuous range of ERK2-GFP expression in the sample population. (**c**) In an overexpression context, inferred fold-changes to 6 kinetic parameters are sufficient to represent signaling dynamics at the population-level with single-cell simulations. Inferred fold changes of the 6 relevant parameters over continuous ERK-GFP expression level are shown. (**d**) Simulated single-cell trajectories based on inferred expression-dependent kinetic functions. Population mean (yellow) and standard deviation (magenta) for data (circle) and simulations (line) are shown. Marginal distribution of data in green; note linear y-axis. (**e**) Marginal distributions of measurement data (green) and single-cell model simulations (orange); note log y-axis. (**f**) Covariance between model components over time for measurements (green circles) and model simulations (orange line). (Inset) Examples of scatter plots comparing single-cell data (green) and single-cell simulations (orange) at given time points. (**g**). Illustrative examples of how the speed of a given reaction (*v*) depends upon a given reaction variable and ERK2-GFP expression. All other reaction components are fixed to visualize the reaction surface. Color corresponds to log of reaction speed *v*.

Experimentally, we used a protein overexpression system based on transient transfection of a GFP-labeled protein of interest (POI) (Lun et al., 2017). By detecting the GFP tag, we measured the amount of non-endogenous protein present in individual cells. Computationally, we modified DISCO to infer continuous functions that describe the kinetic effects of overexpression (Methods, Supplementary Fig. S12). Briefly, the algorithm begins by binning cells according to level of overexpression. The bin-dependent kinetic parameters are then optimized by applying DISCO to each bin while iteratively minimizing the number of kinetic parameters that change as a function of overexpression. For the resultant subset of overexpression-dependent kinetic parameters, a continuous function is used to approximate the overexpression-dependence of each parameter. Finally, the parameters of these expression-dependent functions, which describe changes in kinetic parameters and are not themselves kinetic functions, are fit by DISCO using simulations of the system dynamics.

We previously reported that ERK2-GFP overexpression leads to highly complex responses of MAPK/ERK pathway components to EGF stimulation (Lun et al., 2017; Lun et al., 2019), which thereby provided a challenging test case for DISCO. Transient transfection of ERK2-GFP increased the range of ERK expression by more than two orders of magnitude (Fig. 6b). By using DISCO, we found the huge alterations in signaling responses induced by ERK2-GFP overexpression were surprisingly well represented by overexpression-dependent changes in only six kinetic parameters (Fig. 6c-d). None of these six kinetic parameters, however, changed for very low levels (~0-25th percentile) of ERK2-GFP overexpression, which suggests that cells are robust to a low level of non-endogenous variation in this kinase. Both the range of overexpression at which kinetic changes began and the rate of these changes were parameter-dependent. For example, the parameter *h*_2_, related to dephosphorylation of ppErk, decreased, which suggests the reaction moves towards saturation in the range of the ~025-30th percentile of ERK2-GFP expression. The parameter *g*_6_, related to phosphorylation of Rsk by ppErk, also moves towards saturation (decreased), but only in the ~33-66th percentile of ERK2-GFP expression. Other parameters increased, such as *h*_3_, related to the inactivation rate of pRsk, which suggests the reaction speeds up with increasing ERK2-GFP. Finally, some parameters decreased and then increased, such as the negative feedback parameter *g_FB_*, which might suggest that the parameter changes implicitly capture the effects of otherwise un-modeled or yet-to-be discovered system components.

Signaling systems contain multiple components and may therefore be considered multidimensional surfaces. Having inferred the effects of overexpression on reaction kinetics, we were able to explain the complex single-cell state-space distributions (inset, Fig. 6f) and visualized low dimensional projections of the reaction functions by using fixed values for a subset of reaction variables (Fig. 6g). The results demonstrate how parameters *g_FB_, h*_2_ and *g*_6_ depend non-linearly on the overexpression level of ERK and also on the levels of ppERK. Specifically, this analysis suggests that increasing ERK overexpression decreases *h*_2_ and slows the rate of ppERK inactivation, delaying peak signal of ppERK in cells with high levels of overexpression (Fig. 6d, the tail in the middle inset of Fig. 6f). The overexpression-dependent reduction in ppERK-dependent phosphorylation of RSK and the increase in the rate of the pRSK dephosphorylation reaction (*h*_3_) also explain the circular distribution shape between ppERK and pRSK during signaling (bottom inset of Fig. 6f). Namely, increased levels of overexpression may lead to both higher ppERK levels and lower pRSK levels, but different ranges of overexpression yield both ppERK^*high*^/pRSK^*low*^ and ppERK^*hish*^/pRSK^*intermediate*^ cell states. The model-based inference—that increasing ERK2-GFP inhibits RSK phosphorylation—is supported by previous reports that activation of phosphorylation of p90RSK requires dissociation from inactive ERK (Roux et al., 2003) and that active and inactive ERK compete for binding of p90RSK (Levchenko et al., 2000). It is encouraging that DISCO automatically detects these known facts.

Taken together, DISCO is a powerful tool to explain the effects of protein overexpression in cell samples. Overexpression can have complex kinetic effects, and these effects cannot necessarily be observed in “normal” cells, even with single-cell measurements. Given appropriate measurements of protein expression, however, DISCO provides an effective framework for inferring continuous expression-dependent parameter functions of single-cell states, rather than cell-population states. Thus, one should expect that DISCO can similarly be used to analyze signaling in deregulated or diseased cells.

## Discussion

Up to now, single-cell experiments have been limited because available methods can either measure the dynamics of a few markers or reveal snapshots of many markers, but not both. This trade-off has impeded our understanding of how cell-to-cell variation in expression affects signaling dynamics. Multiplex snapshot single-cell data are becoming more prevalent with improved experimental methods, but analytical approaches to leverage such data for mechanistic modeling remain limited. Here, we present a novel computational approach called DISCO that infers multiplexed signaling trajectories from multiplexed snapshot data. We applied DISCO to study EGF signaling in the MAPK/ERK pathway of HEK293 cells. This analysis revealed that cell-to-cell variation in signaling can be described deterministically, given the initial state of each cell, and that hidden qualitative differences between individual cells can be identified in seemingly homogenous samples. In findings specific to the MAPK/ERK pathway, our model suggests that signal duration is reliably transmitted through the MAPK/ERK cascade, whereas signal amplitude is primarily a fold-change response for each component. A novel auxiliary algorithm permitted inferences of expression-dependent kinetics that are applicable even to extreme cases of overexpression and altered signaling responses. The results demonstrate that DISCO enables the study of cell-to-cell variability in multivariate cellular responses across a broad range of cell states, such as may be observed in disease.

The motivation to develop DISCO derived from a primary limitation of this type of study: Experimental validation of multiplexed single-cell trajectories within individual pathways is not possible. To alleviate this situation, DISCO was developed with minimal assumptions. First, signaling is essentially a deterministic function of cell state. Second, cell-state variables that drive signaling can be measured or inferred. Third, a subset of model parameters can be equal across a cell population. One or more of these assumptions must be violated for the trajectories to become inaccurate. These assumptions actually constitute a subset of those that have been used for model-based inferences of RNA velocities in single cells (La Manno et al., 2018). The assumption of deterministic operation in signaling systems is supported by theoretical arguments regarding the statistics of large numbers of molecules, by simulation studies showing the insignificant contribution of stochastic reaction kinetics to variation in ERK signaling (Filippi et al., 2016; Loos et al., 2018), and by our own results showing that deterministic evolution of individual cell states is sufficient to represent population-level signaling variation, even in an extreme case such as ERK overexpression. The assumption that it is possible to characterize the relevant cellular variation in state or environment can be evaluated by application of multiplexed single-cell cell technologies (Bodenmiller et al., 2012; Giesen et al., 2014; Gut et al., 2018; Chen et al., 2015) of continually increasing dimensionality and, if needed, subsequent subpopulation analyses such as presented here. The assumption that certain kinetic parameters are related to physical properties is a fundamental building block of all biochemical and physiological modeling (Savageau, 1976; Voit, 2013). Thus, where genetic variation in a sample may lead to altered enzymatic activities of a protein, the associated kinetic parameters are genotype dependent.

The primary questions regarding the accuracy of the characteristic signaling response features inferred with DISCO are related to single-cell variation that is missing or mis-attributed in a model. In such a case, we identified at least one subgroup of cells likely to have additional unmeasured (latent) variation in upstream signaling components, which led to signaling behavior that was strikingly different from the other cells in the sample (5). Thus, evaluation of overall model fitness and single-cell response-based analyses in DISCO can identify inaccurate prior assumptions of initial cell states in at least some cases. Our focus on short-term signaling responses simplifies this task because latent variables are expected to be more or less constant over the brief time span of our experiment. As a consequence, sufficient information about latent variables can be inferred from algebraic solutions to steady-state equations. Considerations of much longer time scales, such as those associated with long-term pulsatile dynamics of ERK (Albeck et al., 2013), would necessitate an expanded model and measurements to account for variables no longer in a pseudo steady state.

Single-cell snapshot measurements reveal variations in cell states. An immediate question is how these variations in cell states relate to variations in cell responses. For the MAPK/ERK pathway, we propose that: (1) the signal duration is tightly coupled throughout the pathway in single cells; (2) the variation in signaling amplitudes is much less than variation in the signaling components; and (3) the variation in unmeasured signaling components can be algebraically determined by appropriately analyzing experimental measurements with a deterministic model. The inferences—that the fold-change in ppERK is a population-level response, whereas the duration of ppERK is a single-cell response—are in line with previous imaging-based, information-theoretic studies showing that ERK amplitude can transmit approximately 1-bit of information while consideration of duration can greatly increase information transfer (Cheong et al., 2011; Selimkhanov et al., 2014). Together these results suggest that duration is the informative signal in the ERK pathway and that amplitude primarily serves to raise the signal above noise. Further support is provided by studies that have demonstrated that the duration of ERK signaling is a driver of cell fate (Santos et al., 2007; Ryu et al., 2015).

These results suggest that the interdependent structures of both signaling and expression systems cooperate to reduce variation of cell responses within a population. A clear example is the strong correlation between cell volume, protein abundance and signaling amplitude, which is reasonably explained by isometric scaling of signaling components as cells grow. Thus, signaling systems are structured in such a manner that cells can tolerate the remaining noise and mount appropriate responses in a robust manner. Secondarily, these results suggest caution before unexplained variation in cellular systems is causally attributed to randomness.

Despite the well-coordinated maintenance of signaling response characteristics, this coordination can be impaired in diseases such as cancer, which are associated with mutations and copy number alterations that alter the cellular control of protein expression. Such a system failure was demonstrated with the ERK2-GFP overexpression system. Our analysis revealed that cells operating in abnormal regions of the expression space can have drastically altered signaling response characteristics. Critically in these cases, “normal” cells operated within their physiologically constrained state-space region, whereas “diseased” cells entered new state-space regions in which reaction functions and consequent cellular responses resulted in qualitatively changed behavior. This insight suggests that single-cell data from healthy organisms alone are not sufficient to infer complete representations of reaction kinetics. Instead, cellular systems must be studied both within and outside their normal operating conditions to enable reliable predictions of signaling responses in disease. Such analyses of signaling in deregulated systems will be facilitated by tightly connected experimental and computational analyses, for instance, by combining a protein overexpression system with mass cytometry and DISCO as it is done here.

Here we focused exclusively on signaling, but it is reasonable to expect that the general approach employed in DISCO of combining single-cell simulations, a subset of population-based kinetic parameters, and MMD (or another distribution-free metric) to inform model fitness can similarly be applied to model gene transcription and translation. In these cases, one would extend the relevant time scales, and stochastic simulations and additional associated considerations to account for application-specific uncertainties could be necessary. Finally, DISCO scales linearly with the number of variables, which facilitates its application to data with dimensions on the order of current snapshot cytometry-based measurements, including imaging mass cytometry, where extra measurement dimensions may be used to search for biologically meaningful latent variables.

The data-driven, distribution-free framework developed here represents a qualitatively novel approach to modeling heterogeneity in dynamic biological systems. Combined with the increasing ease of generating multiplexed single-cell snapshot data, we expect the continued development and application of DISCO to deepen our understanding of differential responses in heterogeneous cellular systems.

## Methods

### Cell culture

HEK293T cells, obtained from ATCC, were cultured in DMEM (D5671, SIGMA), supplemented with 10% FBS, 2 mM L-glutamine, 100 U/ml penicillin, and 100 μg/ml streptomycin. For cell passaging or harvesting, cells were incubated with TrypLE Express (Life Technologies) for 2 minutes at 37 °C. Purity and sterility of the cell line were certified by ATCC. Mycoplasma was not detected with the LookOut Mycoplasma PCR Detection Kit (Sigma-Aldrich).

### Quantification of ERK pathway signaling dynamics in response to EGF

Samples were fixed at {0,5,10,15,30,60} minute time points after stimulation with EGF and measured by mass cytometry; the 0-minute sample was unstimulated. Time points were chosen as appropriate to characterize EGF signaling dynamics in the pathway (Lun et al., 2017). Abundances of both total and active proteins were measured for the core pathway components MEK, ERK and p90RSK. In addition, markers for cell volume (GAPDH), cell cycle (IdU, Cyclin B1, pHH3, pCDK1) and cell death (cleaved PARP) were measured using a validated panel of antibodies. Experiments were performed in triplicate for wild type cells, and harvesting, staining and barcoding was performed in parallel. The full antibody panel and staining concentrations are presented in Supplementary Table S4.

### Transfection and stimulation

HEK293T cells were seeded in 6-well plates at a density of 0.7 million cells per well. After 24 hours, cells were transfected with 2 *μ*g ERK2-GFP plasmid (Lun et al., 2017) and 4 *μ*l of jet-PRIME (PolyPlus) per well, with the standard protocol provided by the manufacturer. At 18 hours after transfection, EGF (Peprotech) was added to a final concentration of 100 ng/ml. At 20 minutes before a given EGF stimulation time point, 5-Iodo-2-deoxyuridine (IdU) was added to the medium for a final concentration of 10 *μ*M. At 2 minutes before a given EGF stimulation time point, the medium was replaced by 1× TrypLE to induce cell detachment. At this time point, paraformaldehyde (PFA, from Electron Microscopy Sciences) was added to the cell suspension to a final percentage of 1.6%, and cells were incubated at room temperature for 10 minutes. If EGF stimulation was not necessary in the experiment, cells were directly harvested and crosslinked with PFA. Crosslinked cells were washed twice with cell staining medium (CSM, PBS with 0.5% BSA, 0.02% NaN_3_) and, after centrifugation, ice-cold methanol was used to resuspend the cells, followed by a 10-minute permeabilization on ice or for long-term storage at −80°C.

### Antibody conjugation

The MaxPAR antibody conjugation kit (Fluidigm) was used to generate isotope-labeled antibodies using the manufacturer’s standard protocol. After conjugation, the antibody yield was determined based on absorbance of 280 nm. Candor PBS Antibody Stabilization solution (Candor Bioscience GmbH) was used to dilute antibodies for long-term storage at 4^°^C.

### Barcoding and staining protocol

PFA-crosslinked and methanol-permeabilized cells were washed three times with CSM and once with PBS. Cells were incubated in PBS containing barcoding reagents (102Pd, 104Pd, 105Pd, 106Pd, 108Pd, 110Pd, 113In and 115In) at a final concentration of 100 nM for 30 minutes at room temperature and then washed three times with CSM (Bodenmiller et al., 2012). Barcoded cells were then pooled and stained with the metal-conjugated antibody mix (Supplementary Table S4) at room temperature for 1 hour. The antibody mix was removed by washing cells three times with CSM and once with PBS. For DNA staining, iridium-containing intercalator (Fluidigm) diluted in PBS with 1.6% PFA was incubated overnight with the cells at 4^°^C. On the day of the measurement, the intercalator solution was removed, and cells were washed with CSM, PBS, and ddH_2_O. After the last washing step, cells were resuspended in ddH_2_O and filtered through a 70-*μ*m strainer.

### Mass cytometry analysis

EQ Four Element Calibration Beads (Fluidigm) were added to cell suspensions in a 1:10 ratio (v/v). Samples were analyzed on a Helios mass cytometer (Fluidigm). The manufacturer’s standard operation procedures were used for acquisition at a cell rate of approximately 500 cells per second. After the acquisition, all FCS files from the same barcoded sample were concatenated (Bodenmiller et al., 2012). Data were then normalized, and bead events were removed (Finck et al., 2013) before doublet removal and de-barcoding of cells into their corresponding wells using a doublet-filtering scheme and single-cell deconvolution algorithm (Zunder et al., 2015). Cytobank (http://www.cytobank.org/) was used for additional gating on the DNA channels (191Ir and 193Ir) and 139La/141Pr to remove remaining doublets, debris and contaminating particulates. Dead and mitotic cells were also removed before modeling and analysis. Data were then exported as .fcs files for subsequent analysis.

### Spillover correction

Due to panel design, channel-to-channel spillover as a function of mass resolution (–1 and +1 channels) and oxidation (+16 channels) was only possible in two cases:

1. From channel 143Nd (total ERK) to 144Nd (ppMEK) and 159Tb. (GAPDH)
2. From channel 149Sm (total p90RSK) to 150Sm (total MEK).

Spillover between channels 149Sm and 150Sm was not significant for the measured count ranges and ignored. Due to events with very high counts in 143Nd (from ERK overexpression), spillover from 143Nd was compensated using an estimated 2.2% spillover correction to channels 144Nd (+1) and 159Tb (+16).

### Data normalization and scaling for use in modeling

Experimental measurements were normalized for comparisons across independent experiments and measurements. Before use in modeling, measurement channels were linearly scaled to satisfy biological constraints and facilitate direct physical interpretation of resulting model parameter values. Cells used in fitting and visualization of results were subsampled to reduce unnecessary computational cost. A full description of normalization, scaling and subsampling is presented in the *Supplementary Notes*.

### Maximum mean discrepancy (MMD)

Maximum mean discrepancy (Gretton et al., 2012) (MMD) is a distribution-free method of testing the similarity between samples from two multivariate distributions and has been shown as both computationally efficient and effective when comparing empirical samples from multivariate distributions.

MMD represents the similarity between two distributions (e.g., experimental and simulated single-cell snapshots) as the squared distance between the mean embeddings of distribution features in a reproducing kernel Hilbert space (RKHS). Depending on the kernel *k* associated with the RKHS, infinitely many features may be used to compare the distributions; this feature is in stark contrast to, for example, comparisons of only a few distribution features, such as mean and variance.

Let 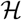 be the unit ball in a RKHS with associated kernel *k*. Given *m* samples from a distribution *X* and *n* samples from a distribution *Y*, an empirical estimate of MMD between *X* and *Y* is

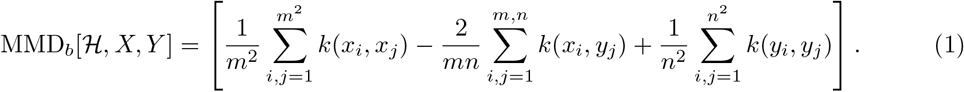

MMD as defined in equation (1) is a biased test statistic, but will still be small if *X = Y* (the true distributions are equal) and large if the distributions are far apart. MMD will be zero if and only if the two samples being compared are equal. Bootstrapping can be used to generate a null distribution of the MMD statistic and determine the statistical significance of the difference between the distributions, but is computationally expensive if performed within an optimization routine. In this work, we are concerned with finding a model that generates distributions similar to experimental measurements, but less concerned with the statistical significance of the model fit to experimental data at any given step of the optimization algorithm (or comparison of alternative models). Thus, it is sufficient for the DISCO procedure to minimize MMD between simulated and experimental distributions, without additional bootstrapping to determine statistical significance. However, we use boot strapping to compare models or parameter sets after optimization, as described below and shown in Supplementary Figures S6 and S5.

### Objective (cost) function for optimization

Given discrete snapshot measurements at times *t* ∈ *T*, corresponding single-cell multiplexed snapshot measurements *D_t_* of *v* variables (e.g., proteins) in *n* cells (D_*t*_ ∈ ℝ^*n×v*^), and a set of *n* continuous single-cell simulated trajectories *Y*(*t*) of the *v* measurement variables, the objective function Cost is defined as:

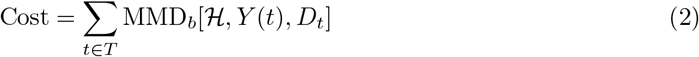

where kernel *k* associated with 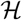 (equation 1) is the Gaussian kernel:

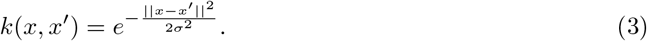

The parameter *σ* scales the width of the Gaussian kernel and was chosen as the median distance between points in the aggregate sample, which is the classical median distance heuristic for selection of kernel bandwidth. For our calculations, the individual observations *Y*(*t*) and *D_t_* were transformed using the natural logarithm of the respective value plus one. The MMD computation was implemented in Matlab (The Mathworks, Inc.) using code available at http://www.gatsby.ucl.ac.uk/~gretton/mmd/mmd.htm.

### Parameter optimization algorithm

Parameter optimization was performed using a combination of global and local search methods. An initial global search was performed using Latin hypercube sampling to select *n* candidate parameter sets from a user input parameter space defined by upper and lower bounds for each parameter.

The cost of each candidate parameter set was evaluated by simulating the single-cell ODE model for a set of *K* cells sampled from the initial time point. Subsequently, the top *m* < *n* candidate parameter sets (those with the lowest cost value) were chosen for local refinement.

Local parameter refinement was performed for each of the top *m* parameter sets using an iterative local approach based upon the Nelder-Mead simplex algorithm (Lagarias et al., 1998) as implemented in `fminsearch` of the Matlab Optimization Toolbox (The MathWorks, Inc.). The iterative approach was used to allow repeated expansion-contraction cycles of the local search region. For iteration *j*, the parameter set output *mj* is input as the initial search point for iteration *j* + 1. In the first local search iteration (j = 1), `fminsearch` is called using a maximum of 50 steps. In all subsequent local search iterations, `fminsearch` is called using a maximum of 150 steps. If, by the end of the second iteration (j=2) the objective function is not less than 98 percent of the initial cost before local refinement, then the local search is terminated. Otherwise, the local search continues to iterate until the cost value for iteration *j* + 1 is not less than 95 percent the cost of iteration *j*.

After local refinement of the top *m* global parameter sets, the single “best” parameter set for use in modeling and analysis was selected. As the cost value for each of the *m* locally optimized parameter sets was calculated using a single subsample of *K* cells for the single-cell model, however, we used bootstrapping to determine the parameter set with the most robust “best” cost. Specifically, after local refinement, we selected the subset of 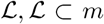 parameter sets with a clear qualitative gap in objective cost compared in the distribution of m cost values. For each parameter set in 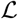, we used bootstrapping to generate independent subsamples of *K* cells for simulation and single-cell model cost evaluation. The single-cell model was not re-optimized and the result is a distribution of model cost values. The parameter set with the lowest mean cost over all bootstrapped cost calculations was taken as the “Best Theta” for use in subsequent model evaluation and analysis.

## Mathematical model of EGF signaling in the MAPK/ERK pathway

The ODE model structure included an input, RAF, MEK, ERK and p90RSK, as well as the known negative feedback loop from ERK to active RAF, as active ERK is known to enhance inactivation of active RAF. The input was modeled as a single bolus corresponding to an addition of EGF. Reaction steps in the MEK/ERK/p90RSK cascade were simplified to include molecular states that were experimentally observable. For example, ERK is activated when it is doubly phosphorylated on threonine 202 and tyrosine 204 (pT202/pY204) by active MEK in a two-step process. However, only the active pT202/pY204 form of ERK was measured, and the transition from inactive to active ERK was therefore modeled as a single step. The initial wild-type (WT) model (Fig. 3) includes 17 total free parameters. 15 parameters are related to kinetics, 1 parameter (*τ*) characterizes the time delay between EGF addition and MEK phosphorylation and 1 parameter (*I_f_*) characterizes any additional residual EGF signaling at the end of the experiment when compared to the initial steady state.

To characterize unmeasured variation across single cells that may affect signaling, we included four single-cell parameters {*k_d_*, *k*_4_, *k*_6_, *k*_8_}: one at each level of the pathway. These parameters were not optimized directly. Rather, the single-cell parameters were algebraically determined by solving the steady-state equation, using a combination of multiplexed measurements at steady state and population-level parameters that were fixed for any given model simulation.

> All models and were implemented in Matlab Release 2015b (The MathWorks, Inc.). Below we include the full mathematical description of the model.

### Model state variables

The model uses eight state variables to describe changes in ERK pathway signaling components. Phosphorylated and total MEK, ERK and p90RSK were measured. After scaling (Methods, *Supplementary Notes*), inactive forms were calculated as Inactive = Total — Active. Total protein levels in each cell were assumed constant over the time scale considered (Supplementary Figure S14). As cells were grown in a mono-layer and EGF was rapidly mixed with medium, the input *I* was assumed to be the same for all cells. The initial value of pRaf was also assumed to be the same for all cells. This choice was made to reduce prior assumptions on the distribution of active RAF. The variation in initial pRAF, as well as of other components upstream of MEK that were not explicitly in the model, were captured by the single-cell parameters *k_d_* and *k*_4_, which were obtained from steady-state measurements (see *Supplementary Notes*).

### Model kinetic parameters

The rate constants *k_d_, k*_4_, *k*_6_ and *k*_8_ were in **Φ**_*k*_ and computed from the steady-state equations as detailed in the *Supplementary Notes*. All other parameters were in **Θ** and, therefore, equal across all cells and used as decision variables in the optimization algorithms.

### Model inputs

The MAPK/ERK pathway components considered in our model were in a pseudo-steady state at time scale of our experiments (less than one hour). Thus, our model must also be at steady state before simulated addition of EGF. We arbitrarily chose the input value of *I*(*t* ≤ 0) = *I_ss_* = 1 as the pre-stimulation steady state input *I* to our model. This value was initially assumed to be the same across all cells in the model (as in Figure 3), but became sub-population-dependent upon the subpopulation analysis in Figure 4. Addition of EGF was simulated as an instantaneous increase in the input to *I*(*t* = 0 + *τ*) = 10. We used a time delay *τ* to represent the delay between experimental addition of EGF to the medium and the time when the signal reached the MAPK/ERK cascade.

In other words, *τ* represents the time it takes for “signal” to be transmitted by reactions, such as receptor-ligand binding, receptor activation, etc., and reach RAF activation. Although the exact value of *τ* for each cell is undoubtedly variable across the population, as not all cells will encounter the EGF signal at the same moment, we assumed each cell to have the same delay (*τ*) to maintain the entirely deterministic nature of our model. Thus, in the model, if time *t* < *τ*, then Input *I* = *I_ss_*. Once time *t* = *τ*, then input is reset to represent EGF addition and Input *I*(*t* = *τ*) = 10. At all other times, input *I* is a dependent variable and determined by solution of the ODE system. The parameter *τ* was a free model parameter, and the inferred value *τ* = 2.02 minutes corresponds well to live cell studies of ERK activation (Ryu et al., 2015). Finally, we found steady-state signaling after addition of EGF was marginally increased compared to pre-stimulation conditions. Thus, for values of *t* > *τ* we used the value *I_ss_* = *I_final_*, which was a free parameter during the optimization (see model equations, below), to limit the depletion of input signal. Expressed differently, before the addition of EGF, the minimum signal can be *I_ss_*, but after addition of EGF the minimum signaling can be *I_f_*.

### Model equations

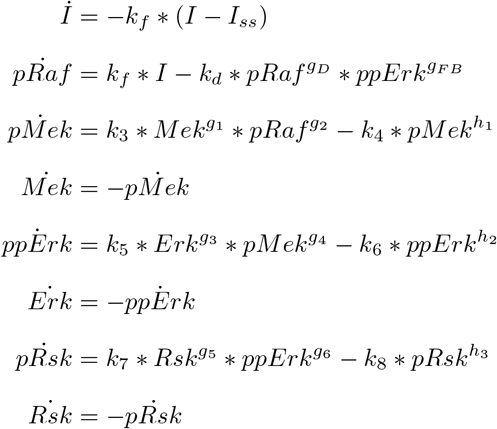

### ODE integration and solver speed-up

The ODEMEX CVode wrapper for Matlab (Vanlier et al., 2012) was used to compile MEX files in C++ using numerical integrators from the SUNDIALS CVode package (Lawrence Livermore National Laboratory, Livermore, CA).

### Model fitting

The 17 free parameters of the model were fit using the global-local optimization approach described above.

Specifically, the WT model was simulated and fit using *K* = 300 single cells randomly subsampled from across the three replicate snapshot mass cytometry measurements per time point during a 60-minute time course of EGF stimulation. For each sampled time point, the six-dimensional cell-state distributions of both total and active forms of MEK, ERK and p90RSK were used to inform model parameters. For the global search step, Latin hypercube sampling was used to generate *n* = 50, 000 samples from the candidate parameter space in Table 4. The top *m* = 50 candidate parameter sets were taken for local refinement. The cost distribution of these 50 global parameter sets after local refinement is shown in Supplementary Figure S5. This algorithm is implemented in `repeatParũptWhile_PowerModel`.

**Table 1:** Model state variables.

| State variable | Description |
| --- | --- |
| $I$ | Input (EGF) |
| $pRaf$ | Active RAF |
| $Mek$ | Inactive MEK |
| $pMek$ | Active MEK |
| $Erk$ | Inactive ERK |
| $ppErk$ | Active ERK |
| $Rsk$ | Inactive p90RSK |
| $pRsk$ | Active p90RSK |

**Table 2:**
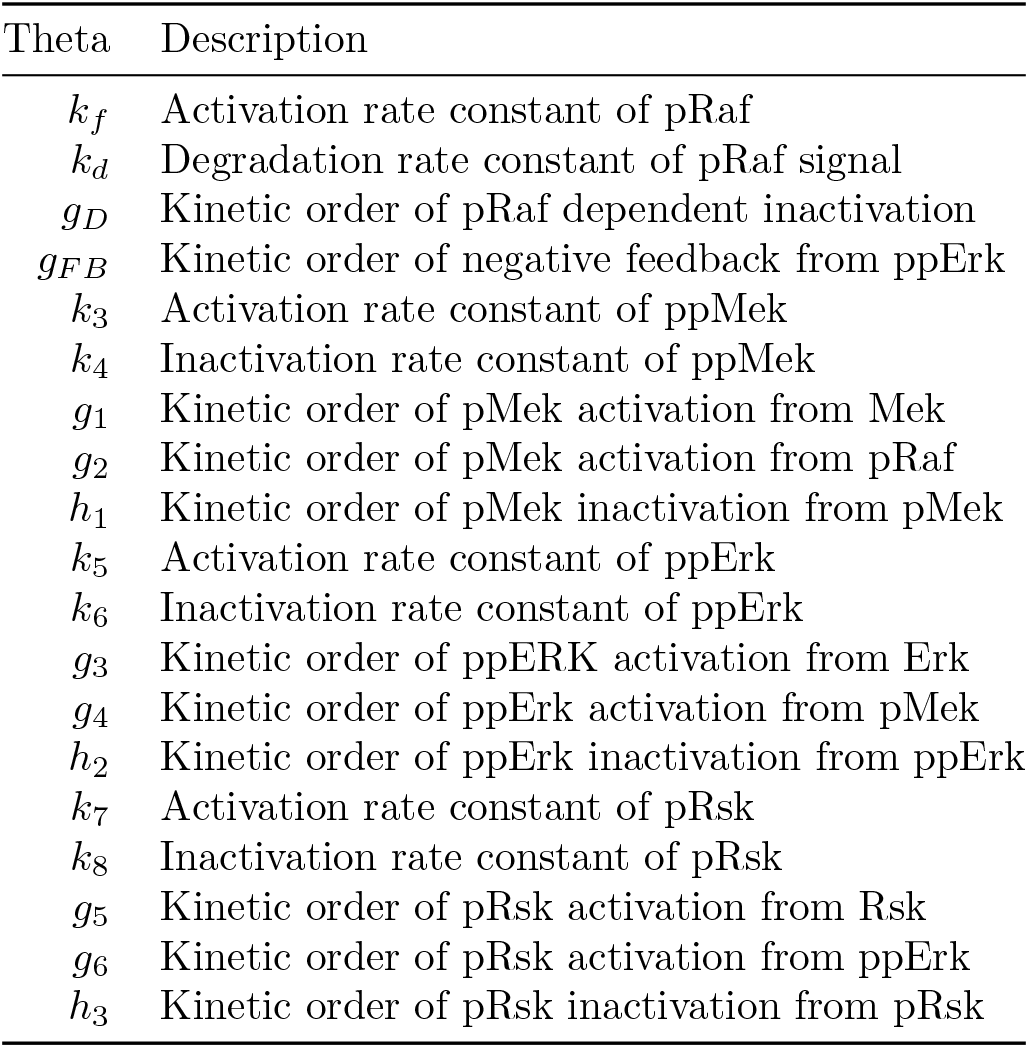
Model kinetic parameters. Note that the parameters {*k_d_,k*_4_,*k*_6_,*k*_8_} ∈ **Φ_*k*_** and therefore computed for each single cell *k* using the steady-state equations.

| Theta | Description |
| --- | --- |
| $k_f$ | Activation rate constant of pRaf |
| $k_d$ | Degradation rate constant of pRaf signal |
| $g_D$ | Kinetic order of pRaf dependent inactivation |
| $g_{FB}$ | Kinetic order of negative feedback from ppErk |
| $k_3$ | Activation rate constant of ppMek |
| $k_4$ | Inactivation rate constant of ppMek |
| $g_1$ | Kinetic order of pMek activation from Mek |
| $g_2$ | Kinetic order of pMek activation from pRaf |
| $h_1$ | Kinetic order of pMek inactivation from pMek |
| $k_5$ | Activation rate constant of ppErk |
| $k_6$ | Inactivation rate constant of ppErk |
| $g_3$ | Kinetic order of ppERK activation from Erk |
| $g_4$ | Kinetic order of ppErk activation from pMek |
| $h_2$ | Kinetic order of ppErk inactivation from ppErk |
| $k_7$ | Activation rate constant of pRsk |
| $k_8$ | Inactivation rate constant of pRsk |
| $g_5$ | Kinetic order of pRsk activation from Rsk |
| $g_6$ | Kinetic order of pRsk activation from ppErk |
| $h_3$ | Kinetic order of pRsk inactivation from pRsk |

**Table 3:** Model inputs.

| Input | Description |
| --- | --- |
| $I$ | Input (EGF) |
| $I_{ss}$ | Input at steady state (before EGF addition) |
| $I_f$ | Minimal input level after EGF addition |
| $\tau$ | Time delay between addition of EGF and initial pRaf signaling |

**Table 4:**
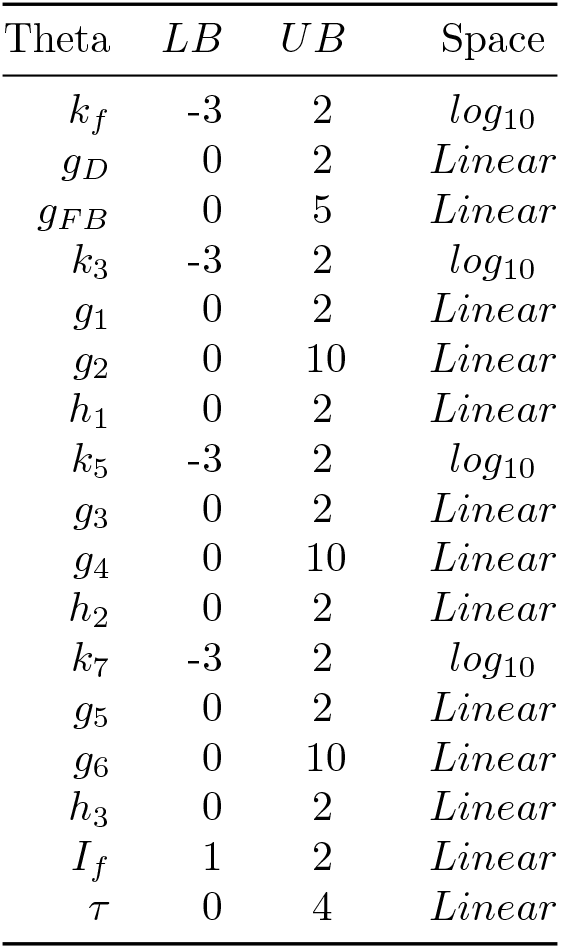
Lower (LB) and upper (UB) bounds and associated space for initial Latin hypercube sampling of each parameter during global search step. Note the local search step was unconstrained and could therefore search beyond these bounds.

| Theta | LB | UB | Space |
| --- | --- | --- | --- |
| $k_f$ | -3 | 2 | $\log_{10}$ |
| $g_D$ | 0 | 2 | *Linear* |
| $g_{FB}$ | 0 | 5 | *Linear* |
| $k_3$ | -3 | 2 | $\log_{10}$ |
| $g_1$ | 0 | 2 | *Linear* |
| $g_2$ | 0 | 10 | *Linear* |
| $h_1$ | 0 | 2 | *Linear* |
| $k_5$ | -3 | 2 | $\log_{10}$ |
| $g_3$ | 0 | 2 | *Linear* |
| $g_4$ | 0 | 10 | *Linear* |
| $h_2$ | 0 | 2 | *Linear* |
| $k_7$ | -3 | 2 | $\log_{10}$ |
| $g_5$ | 0 | 2 | *Linear* |
| $g_6$ | 0 | 10 | *Linear* |
| $h_3$ | 0 | 2 | *Linear* |
| $I_f$ | 1 | 2 | *Linear* |
| $\tau$ | 0 | 4 | *Linear* |

After local refinement, we selected the subset parameter sets below a clear qualitative gap in objective cost compared to the distribution of *m* = 50 cost values. This selection was calculated using a gap in cost values of at least 0.5 (Supplementary Fig. S5, implemented in `optimizationCostDistributionThreshold`). We note that this cost width gap is qualitative and the threshold criteria were tuned depending on the expected distribution. For each parameter set, we used bootstrapping to select 500 different cell subsamples (of K = 300 cells) for simulation and model cost comparison (implemented in the function `ThetaCompare`). The parameter set with the lowest mean over all 500 cost calculations was taken as the “Best Theta” (Θ*) and used in subsequent model evaluations and analyses (Supplementary Fig. S5).

### Bootstrapped evaluation of model fitness

To better estimate the quality of any single-cell model fit to experimental data, we used bootstrapping. To estimate the MMD distribution of the single-cell model, we generated *J* = 500 sets of *K* = 300 cells each by random sampling without replacement from across replicates for each of the *T* = 6 experimental time points. Then, we simulated the single-cell model using the *j^th^* sample set and calculated the associated MMD cost distribution MMD_*j,t*_ at each time point *t*. The total MMD cost of the model for the *j^th^* cell subset was calculated as

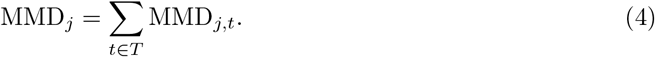

To estimate the variation in MMD due to experimental variation (harvesting, staining, barcoding), we generated *Ĵ* = 500 samples of *K* = 300 cells from each of the *T* = 6 experimental time points for each of the three experimental replicates. We then calculated a null MMD distribution at each (*t* ∈ *T* = 6) time point 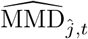 by comparing each time point across the 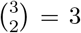 unique paired cross-replicate comparisons for all *Ĵ* = 500 samples of *K* = 300 cells each. The total model MMD cost for a given cross-replicate comparison was calculated as

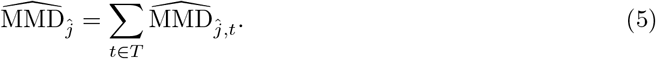

We then compared the bootstrapped model distribution ∪_*j∈J*_ MMD_*j*_ to the null distribution 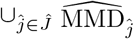, which was calculated using experimental replicates (fig. 3b). For this comparison, we omitted the initial time point *t* = 0 because, by definition, the MMD for the model at the initial time point is equal to zero (we are comparing the exact same distribution). Comparison of bootstrapped model and null MMD distributions by time are shown in Supplementary Figure S6. This analysis is implemented in `ThetaCompare`.

### Calculation of signaling response features

For each signaling phosphoprotein *X_i_* in cell *k* during a time period *T*, the following signaling response features were calculated:

Amplitude: the maximal increase in protein phosphorylation relative to initial cell state:

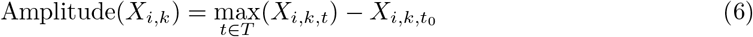

Duration: the time period during which protein phosphorylation is above an amplitude threshold *α*. For our analysis, we set *α* = 0.75:

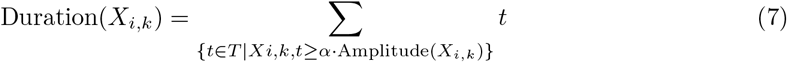

Fold-amplitude: the amplitude normalized (divided) by initial state:

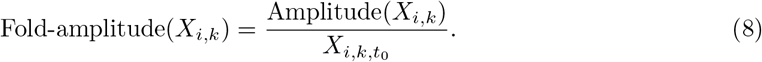

Peak-time: the time of peak amplitude:

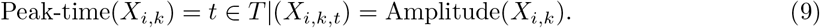

Number of peaks (nPeaks): number of peaks (first time derivative = 0, second time derivative < 0). This function was implemented using the Matlab function `findpeaks` with the additional criterion that a peak must have a prominence greater than 10 percent of peak amplitude for state *X_i_* in cell *k*.

Decay rate (Decay): the time from peak amplitude until the phosphoprotein signal drops below a fractional threshold *α* for peak amplitude. For our analysis, we set *α* = 0.75:

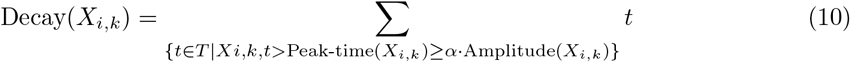

Integral: the sum over all (phosphoprotein) signal above the steady-state level over the entire time course:

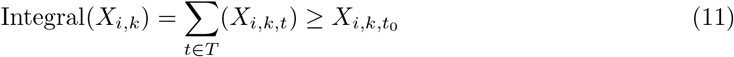

These features were calculated for each cell. Similarly, the measured state variables and inferred single-cell parameters were calculated as shown in `Figure_3_Subpopulation_Analysis` and included in the Supplementary Files (`CellFeatures.mat`). For the analysis in Figure 5, these features were calculated on cells with subpopulation-dependent steady-state inputs (see below); they are included in Supplementary Files (`CellFeaturesSubpop.csv`). As many of these features are correlated, we focused the analysis presented here on the amplitude, duration and fold-amplitude.

### Mahalanobis distance to calculate population outliers

The Mahalanobis distance (*D_M_*) provides a measure of the distance between a point and a distribution (Mahalanobis, 1936). Intuitively, *D_M_* calculates the Euclidean distance of the point from the mean (the center of mass) of the distribution, accounting for the covariance structure (the direction of spread) of the distribution. Given two points with equal Euclidean distance from the distribution center, the point that lays more in line with the distribution spread will be judged as “relatively closer” and have a lower Mahalanobis distance.

Given a point represented by a vector 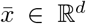 and a distribution with mean ***μ*** ∈ ℝ^*d*^ and covariance matrix *S* ∈ ℝ^*d*^, the Mahalanobis distance *D_M_* of 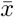 from the distribution is defined as:

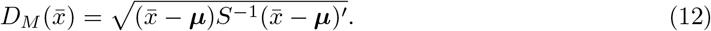

The Mahalanobis distance distribution for the measurement space was calculated using total and phospho-MEK, ERK and p90RSK at steady state. The Mahalanobis distance distribution for the response space was calculated using the amplitude and duration values of phospho-MEK, ERK and p90RSK. The outlier cutoff (as shown in Fig. 4b) was taken as the absolute value of the second percentile of the respective Mahalanobis distribution. The Matlab function mahal calculates Mahalanobis distance.

### Gaussian mixture model for subpopulation clustering

To examine potential subpopulations in the feature space of our cells, we use a regularized Gaussian mixture model using the function `fitgmdist` implemented in Matlab. A regularization value of 0.1 was chosen as sufficient to penalize assignments of small populations while being in a relative robust range. The total number of subpopulations was chosen using the Akaike information criterion (AIC) (Akaike, 1974). Cells were assigned to clusters according to the maximum posterior probabilities (i.e., hard clustering).

In the response space, we considered the amplitude and duration of pMek, ppErk and pRsk. Cell assignment according to posterior probabilities and the number of cells per cluster are shown in Supplementary Figure S7.

To compare response-based clusters to potential measurement-based clusters, we applied the same approach to measurement-space defined by total and phospho-MEK, ERK and p90RSK. Supplementary Figure S7 compares the response- and feature-based clusters in the relative initial activation space (as in Fig. 4e). Supplementary Figure S8 illustrates the response- and feature-based clusters across the full combined feature space.

This analysis is implemented in `Figure_3_Subpopulation_Analysis`.

### Subpopulation-dependent optimization of steady-state input

To determine response-based subpopulation-dependent parameter values for the steady-state input values, the steady-state input was taken as an additional model parameter. Thus, all cells assigned to a subpopulation *p* were given the same steady-state input level *I_ss_p__*. The values of *I_ss_p__* (a total of five parameters, one for each response-based subpopulation) were optimized following the global-local optimization algorithm described above. Latin hypercube sampling was used to generate 500 initial search points in the range of 0.1 to 10 across all five subpopulation parameters. This search range represented the original steady-state input value of one +/− an order of magnitude. The top 12 global parameter sets were taken for local refinement. The initial threshold for cost improvement was a final cost value less than 99 percent of the original after 100 optimization search steps. Subsequently, the same 99 percent threshold for improvement was applied after 200 steps. The algorithm ended when the threshold was not met (see implementation in `repeatParũptWhile_PowerModel_SubPop`).

### Correlation analysis of cell features

Correlation analysis of single-cell measurement and response features, as well as the single-cell variation *k_d_, k*_4_, *k*_6_, *k*_8_ determined by steady-state equations, was performed using Spearman’s rank correlation.

First, partial least squares regression was applied to remove confounding effects of variation in cell size as measured by GAPDH. Next, the Spearman’s rank correlation between features *X* and *Y* was calculated:

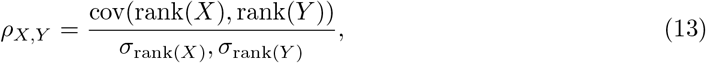

where the function rank(o) provides the relative rank of observations of a given feature. We used rank, rather than linear (e.g., Pearson’s *r*) correlation for analysis due to the nonlinear nature of the system and associated response features. To represent the relative power of the rank correlation in one feature to explain variation in the rank of another variable, we used the signed squared rank correlation sgn(*ρ*) · *ρ*^2^.

This analysis is included in the R script `Figure_4_CorrelationPlot`.

### Coefficient of variation of cell features

The coefficient of variation (CV) represents a normalized measure of distribution width. For a distribution with mean *μ* and standard deviation *σ*, the CV is the given by

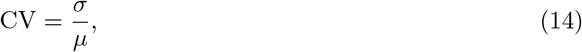

i.e., the standard deviation is normalized by the mean.

This analysis is included in the R script `Figure_4_CorrelationPlot`.

### Data and parameter normalization for modeling ERK-GFP overexpression

The WT and overexpression experiments were performed and measured far apart in time. To account for differences in staining and sensitivity, measurements from the overexpression experiments were scaled to WT measurements based on steady state. To account for changes in instrument sensitivity, which led to slight variation in the signal range for different variables, model parameters were locally re-optimized.

Specifically, overexpression experiments included FLAG-GFP overexpression samples (controls) that were stained, barcoded and measured in conjunction with ERK2-GFP overexpression samples. Scaling factors for each protein were calculated by normalizing the average steady-state values in FLAG-GFP samples to the mean steady-state value across WT replicates. The scaling factors were then applied to the FLAG-GFP control and ERK2-GFP samples across all time points (Supplementary Fig. S10). Next, a local optimization search was performed using `fminsearch`, starting with the WT model parameter set, to fit the FLAG-GFP data. Supplementary Figure S11 illustrates the model compared to the FLAG-GFP data before and after fitting. For the subsequent analysis of the effects of ERK2-GFP overexpression on signaling kinetics, the FLAG-GFP control parameter set was taken as wild type.

### Inferring overexpression-dependent kinetic parameters and associated continuous functions

To find a minimal subset of model parameters that should change as a function of overexpression, we developed an extended algorithm based upon DISCO (Supplementary Fig. S12). This algorithm proceeds in the following steps:

1. Bin data by overexpression level (POI-GFP or total protein, for example).
2. Select a subset of bin-dependent model parameter values (those model parameters that may depend on overexpression).
3. For each bin, re-optimize all bin-dependent model parameters to generate bin-dependent parameter values.
4. For each bin-dependent parameter, calculate the decrease in model fitness that results from fixing the parameter across bins (e.g., how sensitive is the model to removing this parameter from the overexpression-dependent set).
5. Remove those parameters from the overexpression-dependent set that can be fixed without decreasing model fitness (increasing the cost function) beyond a threshold.
6. If parameters were removed, return to step 2 and repeat, otherwise, continue.
7. Fit the bin-dependent changes in model parameters, which may be considered piecewise linear functions, with candidate continuous functions (e.g., a 3^*rd*^ degree polynomial and a Hill function). Single-cell expression-dependent parameter values may be estimated using a linear interpolation of the overexpression-bin-dependent values. Fitting of continuous functions may then be weighted by cell density and the MMD cost of each bin.
8. Select the continuous functional form for each expression-dependent parameter with the lowest *R*^2^ value.
9. To determine how well the continuous functions represent expression-dependent parameter changes in the context of the model, simulate the single-cell model using the selected continuous functions to capture expression-dependent model parameter values and calculate the associated MMD cost.
10. As the continuous functional form that best fits the binned parameter values may not provide the best fit for the dynamic model, hierarchically compare the use of alternate functional forms for each expression-dependent parameter when simulating the single-cell model. This hierarchical comparison greatly reduces the necessary number of combinatorial comparisons.
11. If fitness of the dynamic single-cell model improves from a switch in functional form, switch the functional form and apply step 10 for the next parameter. Otherwise, apply step 10 for the next parameter.
12. Once the functional forms to describe expression-dependent model parameters have been selected, refine the parameters of the continuous functions (that represent expression-dependent model parameters) using DISCO applied to the overexpression data. We emphasize that all WT model parameters remain fixed: only the parameters for the functions that describe expression-dependent changes to model parameters are being fit.

For our specific implementation of this algorithm, we used 8 equally spaced bins of ERK2-GFP in the log(1+GFP) space (Supplementary Fig. S12). For bin-dependent parameter search using DISCO, 125 cells were sampled without replacement from each bin. The choice of bin number was made to ensure sufficient cells could be sampled per bin. We assumed the model parameters for the final steady-state input *I_f_* and the input delay *τ* should not be expression dependent. The remaining 15 model parameters were allowed to depend on ERK2-GFP expression. Serial removal of parameters as described in steps 2-5 was based on the sensitivity threshold:

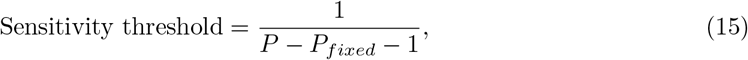

where *P* equals the total number of WT model parameters and *P_fixed_* the number of WT model parameters assumed to be independent of expression level. Thus, we used the sensitivity threshold 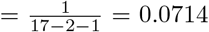 (or a ~ 7% decrease in model fitness), which resulted in three total rounds of refinement (Supplementary Fig. S12). When applying DISCO and local search to find expressionbin-dependent parameter values, we used a serial approach to improve the smoothness of parameter value changes across bins (Supplementary Fig. S12): the initial search point of bin *b* was the output of bin *b* — 1. For example, the WT parameter values were taken as the initial point of the optimization for the first bin; for the optimization of parameter values associated with the second bin, the output from the optimization of the first bin was taken as the initial point; and so on. Based on the expression-dependent parameter changes across bins, we tested use of both 3^*rd*^ degree polynomials and sigmoidal Hill-type continuous functions as candidate descriptors (Supplementary Fig. S12). The intercept for both functions was fixed at the WT model parameter value, which resulted in three free parameters for each function. In the event that use of one functional form caused failure in the solution of the ODE solvers for the single-cell model, the alternate functional form was used.

The algorithm to find the minimum subset of expression-dependent parameters is implemented in `findMinSetBinnedTheta`. The algorithm to select and fit continuous functions to the expression dependent parameter changes is implemented in `findContParams`.

## Figures

For each of the main figures, we included the associated data, code and README files with specific instructions to reproduce the associated analyses, figure and supplementary figure panels. Where random sampling is performed, exact results will depend on the random seed.

## Data availability

The datasets generated during and/or analysed during the current study will be available online upon publication.

## Code availability

All relevant code is included in the supplementary materials.

## Acknowledgements

The work of JDW was supported by the G.F. Amelio Fellowship awarded by the Georgia Institute of Technology College of Engineering, a National Science Foundation Graduate Research Fellowship under Grant No. DGE-1148903 and a Whitaker International Fellowship awarded by the Institute of International Education. This work was also supported by (NSF/MCB 1517588 (PI: EOV) and NSF/MCB (PI: 1411672, PI: D. Downs)). BB’s research is funded by an SNSF R’Equip grant, an SNSF Assistant Professorship grant (PP00P3-144874), by the European Research Council (ERC) under the European Union’s Seventh Framework Program (FP/2007-2013)/ERC Grant Agreement n. 336921 and an NIH grant (UC4 DK108132). The authors would like to thank Vito Zanotelli for recurring discussion, and Natalie de Souza and Will Macnair for their thoughtful discussions and comments on the manuscript. Will Macnair is credited with the dubbing of “DISCO.”

## Author contributions

JDW and EOV conceptualized DISCO. JDW designed and performed the modeling and analyses. JDW, X-KL and BB designed the experiments. X-KL and JDW performed the experiments. JDW, EOV and BB wrote the manuscript. All authors reviewed and edited the final manuscript.

## Additional information

The authors declare no competing financial interests.

## Supplementary Notes

### Introduction

Here, we provide additional details on *Distribution Independent Single-Cell ODE modeling* (DISCO) as presented in the main text. This material includes comparison of DISCO to other modeling approaches based on parametric distributions or the population average, detailed description of data scaling and exclusion for use in modeling, calculation of parameter profiles for the WT model presented in the main text and, finally, an example of how steady-state equations are used to infer latent cell-to-cell variation at different points in a model.

#### Summary of modeling approaches to represent cell-to-cell variation

**Table S1:**
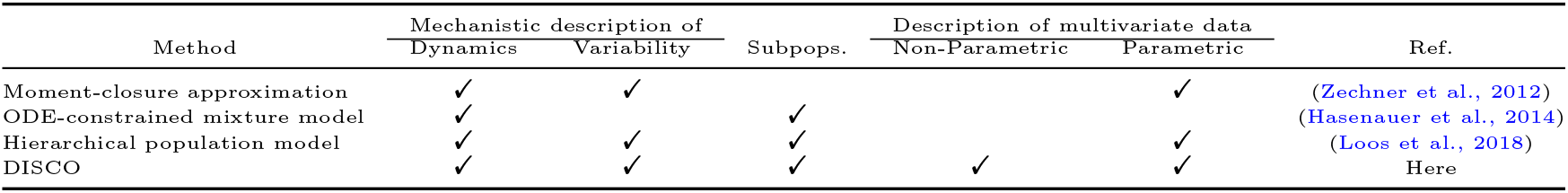
Summary of methods to represent cell-to-cell variation in dynamics.

| Method | Mechanistic description of |  | Subpops. | Description of multivariate data |  | Ref. |
| --- | --- | --- | --- | --- | --- | --- |
|  | Dynamics | Variability |  | Non-Parametric | Parametric |  |
| Moment-closure approximation | ✓ | ✓ |  |  | ✓ | (Zechner et al., 2012) |
| ODE-constrained mixture model | ✓ |  | ✓ |  |  | (Hasenauer et al., 2014) |
| Hierarchical population model | ✓ | ✓ | ✓ |  | ✓ | (Loos et al., 2018) |
| DISCO | ✓ | ✓ | ✓ | ✓ | ✓ | Here |

#### Application of DISCO to benchmark problem for inference of latent subpopulations

To compare DISCO to modeling approaches based on parametric distributions representing cell-to-cell variation in signaling, we used as a benchmark a problem documented in (Hasenauer et al., 2014) and (Loos et al., 2018). The problem was designed to test the ability of methods to infer latent cell populations and the associated kinetic, mixing and distribution parameters, which is not the same goal as DISCO *per se* as DISCO is not constructed to rely on parametric distributions. Specifically, the problem is a conversion reaction with three kinetic parameters and two cell populations (Supp. Fig. S1a). There are three kinetic parameters {*k*_1*p_i_*_,*k*_2_,*k*_3_} where *p_i_* ∈ {*p*_4_,*p*_2_} characterizes the dynamics of each subpopulation *p*_1_ or *p*_2_. Parameter *k*_2_ is constant across all cells. Parameter *k*_3_ is a single-cell parameter drawn for a log-normal distributions with mean *β*_*k*_3__ and standard deviation *σ*_*k*_3__. The proportion of cells in each population *p*_1_ or *p*_2_ is represented by the parameter 0 ≤ *w* ≤ 1 such that the fraction of cells in *p*_1_ = *w* and *p*_2_ = 1 — *w*.

To determine a ground truth for comparisons, single cells are sampled and single-cell trajectories are simulated according to the chosen parameter values. At the given time points, single-cell observations of the state variable *X*_2_ are made. To simulate experimental error, multiplicative Gaussian noise is added according to the parameter *σ_noise_*. For our problem, the cell labels for each time point where then permuted to simulate observation of independent experimental samples at each time point.

We applied DISCO to this problem, but with the altered goal (of the DISCO approach; the goal is rather defined by the previous studies) of inferring the cell population mean *β*_*k*_3__ and standard deviation *σ*_*k*_3__ parameters, rather than single-cell kinetic parameters. Supplementary Figure S1b illustrates the model fit. We used Monte Carlo sampling of 10,000 points around the inferred parameter set to generate parameter confidence intervals (Supp. Fig. S1c). The true parameter values fell within the 99 percent confidence interval for all parameters except for *σ_noise_*. This is due to the construction of DISCO-based modeling: unmeasured single-cell variation is attributed to single-cell parameters, *k*_3_ in this case, using steady-state equations. This is illustrated by the inference error between *σ*_*k*_3__ and *σ_noise_*, where the results show that the missing variation in measurement noise is attributed to single-cell variation in k3. Intrinsically, that is because these two values are only separable in a single-cell construction such as DISCO with additional observations to characterize experimental variation. The parameters to characterize the population mean of the single-cell parameter *β*_*k*_3__ and population mixing component w were near perfectly identified.

We compared signal-cell simulations to the known grown truth, which showed that a number of individual cells were mis-assigned to the opposite population (Supp. Fig. S1d). This is to be expected without additional information on cell state and was a primary motivation for the development of DISCO, which is designed to be used with increasingly multiplexed measurements that, for example, can be used to inform the population assignment. Supplementary Figure S1e illustrates the model results under the assumption that additional information may be used to inform the decision of whether a cell belongs to population 1 or 2. In total, these results illustrate the ability of DISCO to identify latent populations in a benchmark test designed for use with a parametric distribution, rather than an explicit single-cell, modeling approach.

#### Comparison of DISCO to classical population modeling based on means

Although we illustrated several advantages of DISCO compared to classical population-average modeling approaches, such as the ability to infer latent cell populations (Fig. 4), we also applied a population-average modeling approach to our data for comparitive purposes. We applied the same global-local optimization approach to simulate and fit the population average by minimizing the root mean squared error (RMSE) between model simulation and measured data summed over all measurement time points. Not surprisingly, as a different optimization task, different kinetic parameter values were identified (see blue dots in Supplementary Figure S4). Simulation of the population mean using these parameter values resulted in highly underdamped oscillations (Supp. Fig. S3), which is inconsistent with the established signaling behavior of the pathway and suggests that the population-average model is less constrained by the data compared to the single-cell model. This analysis further illustrates how DISCO leverages additional information in multivariate singlecell data and outperforms the classical population-average modeling approach.

**Figure S1:**
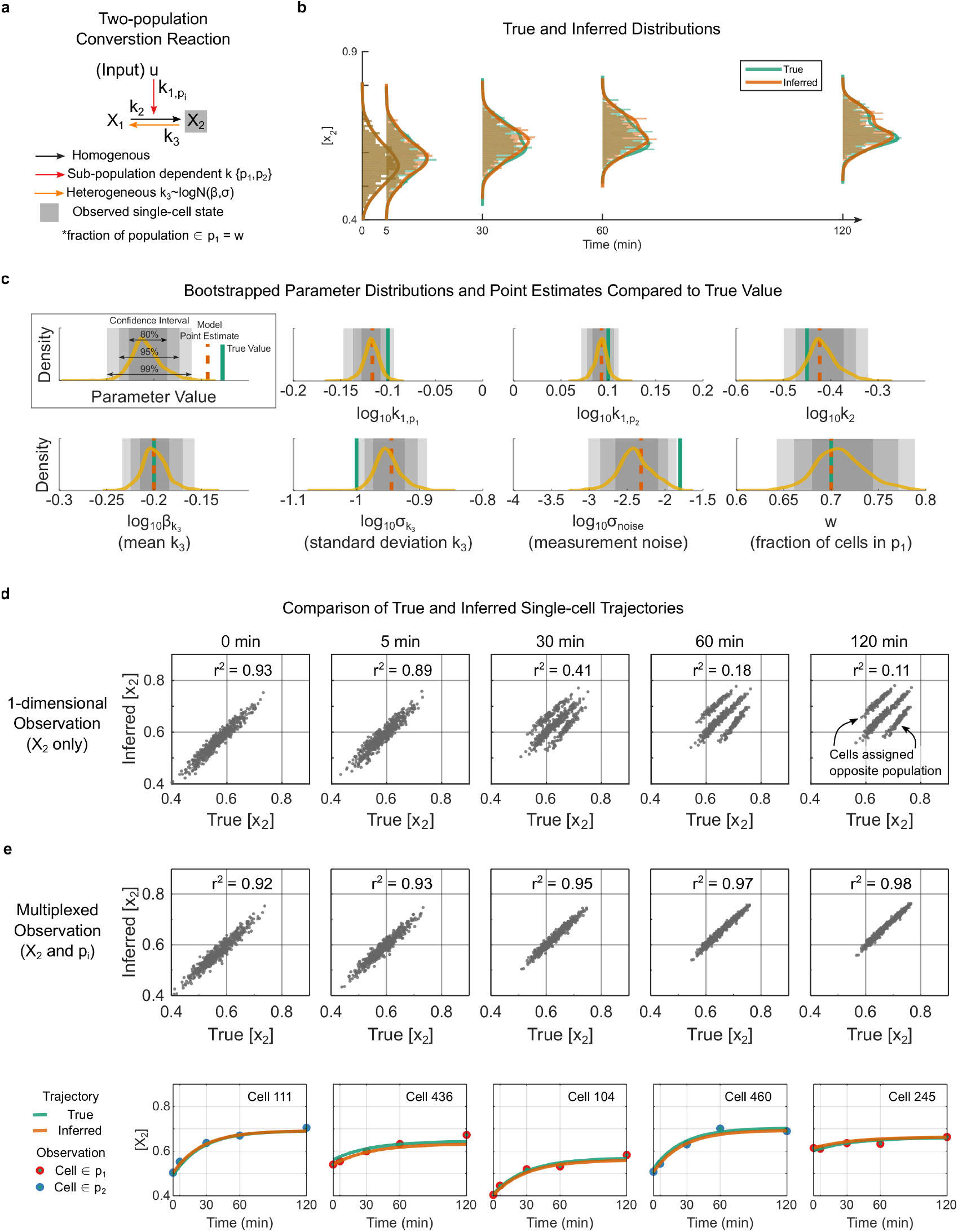
Application of DISCO to the simulated two-population benchmark problem from (Loos et al., 2018) and (Hasenauer et al., 2014). (**a**) Reaction model annotated with state variables and kinetic parameters. Kinetic parameters can be homogeneous (constant for a population), subpopulation-dependent or single-cell-dependent. (**b**) Comparison of true and simulated (based on inferred parameters) marginal distributions. (**c**) True parameter values (green line), inferred point estimates (orange dashed line) distributions (yellow) and associated confidence intervals (gray bars). (**d**) Comparison of simulated single-cell trajectories to known true trajectories. (**e**) Comparison of simulated single-cell trajectories to known true trajectories, exploiting additional information on population assignment. (Bottom) True (green line) and inferred (orange line) single-cell trajectories based on noisy snapshot measurements (dots) for 5 randomly selected cells.

**Figure S2:**
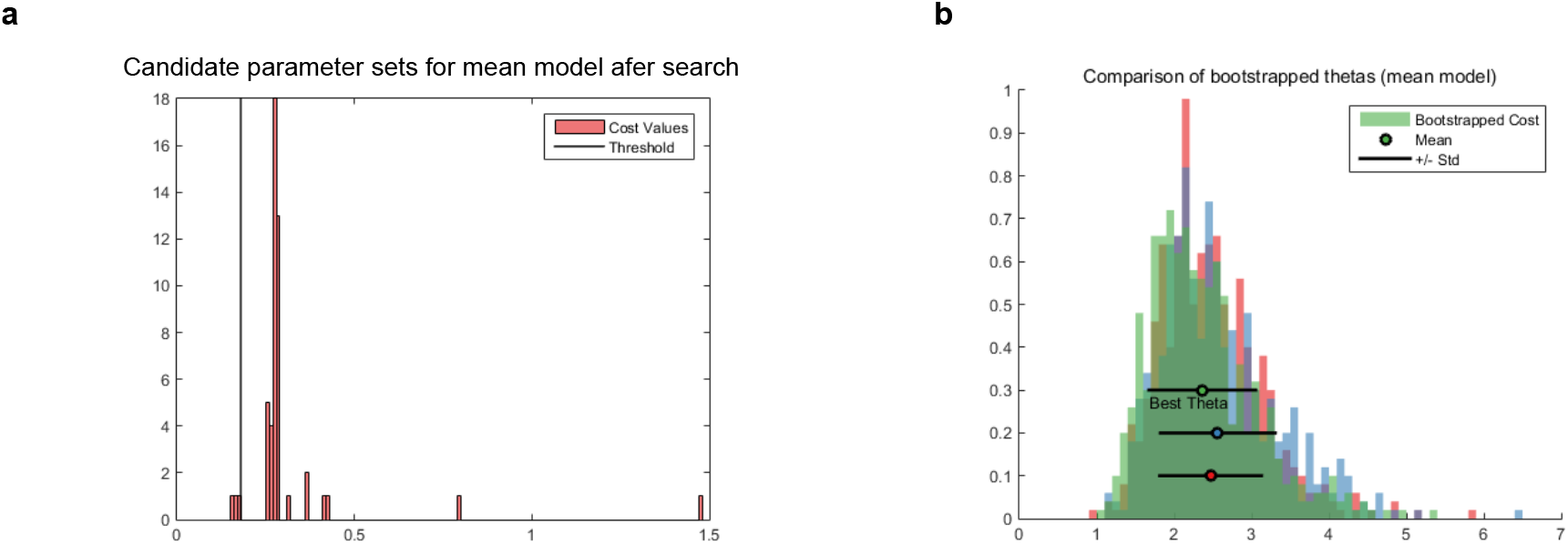
Selection of mean model parameter set. (**a**) Distribution of WT model cost (RMSE) for the top 50 (out of 50,000 initial parameter sets) after local search refinement. Parameter sets with model cost less than the threshold were selected for bootstrapped evaluation of robustness. (**b**) Distribution of WT model costs based on repeated sampling (without replacement) and simulation of mean of cell subsets for the parameter sets below the threshold in panel (a). Shown are the mean (dot) +/− standard deviation (line) for each candidate parameter set. The parameter set labeled “Best Theta” was selected for subsequent model analysis.

**Figure S3:**
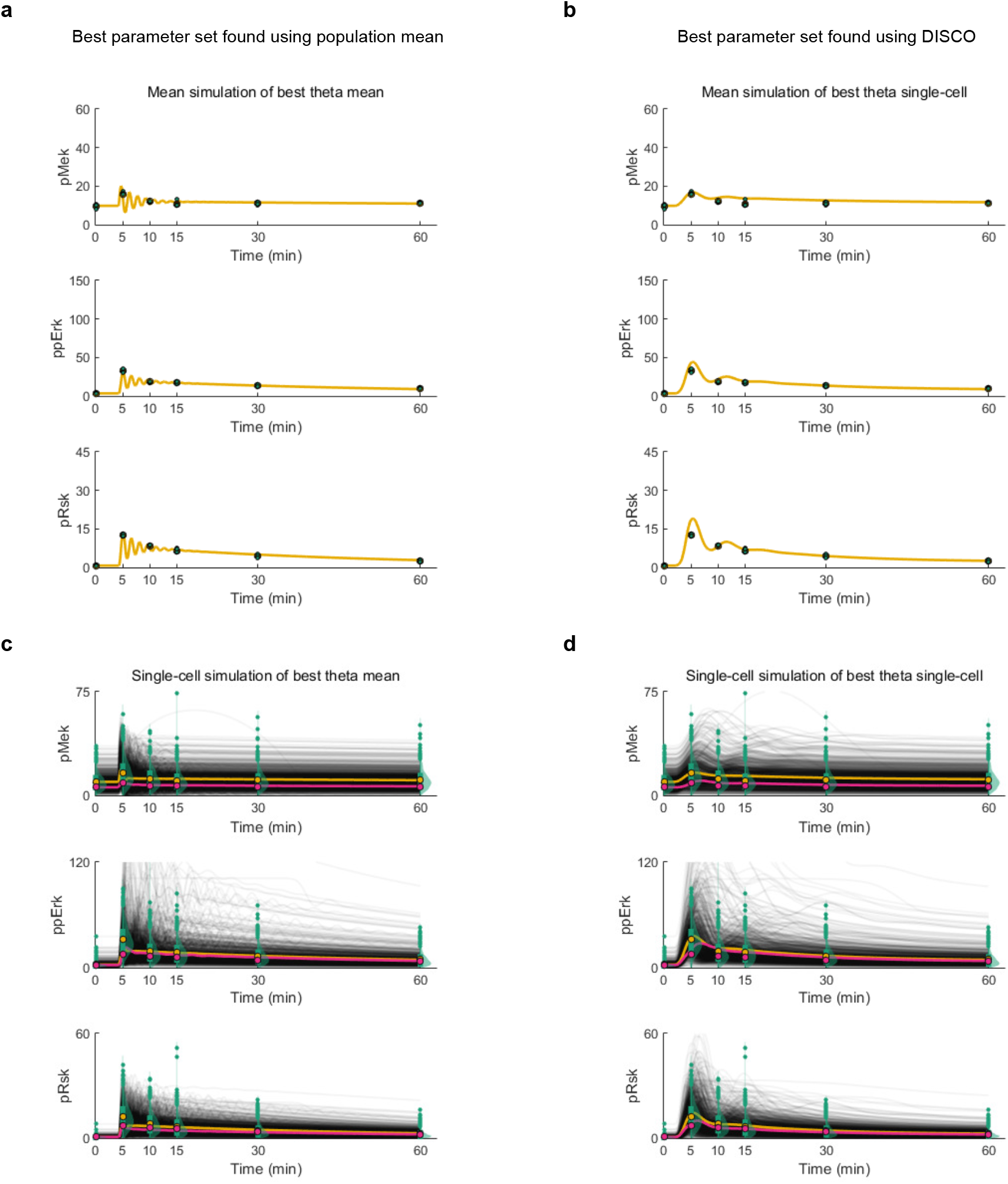
Comparison of DISCO to population-mean model (**a**) Simulation of the population mean using the parameter set found with the population-mean modeling approach. (**b**) Simulation of the population mean using the WT parameter set identified with the single-cell approach in DISCO (the same parameter set as in the main text). (**c**) Use of the best population-mean parameter set to simulate single cells. (**d**) Use of the best DISCO parameter set to simulate single cells.

#### Data normalization and scaling for use in modeling

DISCO uses single-cell measurement values to represent different ODE model instances. This approach is advantageous because it is simple; however, it requires attention to errors in individual cell measurements and, in the case of mass cytometry, rescaling and normalization of measurement channels for use in modeling.

##### Single-cell exclusion criteria and resampling

Antibody labeling and cell staining were optimized to minimize the number of cell events with zero or low ion counts in measurement channels used for modeling (e.g., total ERK). Furthermore, measured cell events with fewer than 5 ion counts in a subset of “filter” channels were excluded. The filter channels were total MEK, total ERK, total p90RSK and GAPDH. The choice of five counts was made as a trade-off between the number of discarded events, which increases as the threshold increases, and the uncertainty of single-cell measurements, which is a decreasing function of ion counts. Such thresholding may introduce bias across samples if, for example, the active form of a protein is on average low in abundance at one time point, but increases in abundance upon stimulation. In this case, thresholding may remove many “low” cells from the first time point that ultimately should represent some of the cells at later time point.

In ERK overexpression experiments, the number of ion counts measured could exceed the linear response range of the mass cytometer. As nonlinearities in measurements violate the optimization objective function as implemented, we excluded cell events with ion counts greater than 10,000 before data sampling and modeling.

##### Data scaling

Mass cytometry provides relative values of protein abundance. Absolute values depend on factors such as antibody labeling efficiency, antibody staining and detection sensitivity. While relative differences between measured protein abundances can often be subsumed into reaction parameters in a model, we rescaled measurements to physiological values.

Average protein abundance values in HEK293T cells were obtained from the MaxQB database (http://maxqb.biochem.mpg.de/mxdb/) (Geiger et al., 2012). The average of iBAQ abundance values across the three experimental replicates was used to determine average abundance (Supplementary Table S2). In cases where an antibody detected multiple isoforms of a protein (e.g., ERK1 and ERK2) the average abundances of each isoform were summed. Abundance values were scaled to concentrations using the average HEK cell volume (Boss et al., 2013) of 1996 *μ*m^3^. These values were used to estimate the average concentration 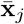 of each protein *j* in a cell. Protein measurements were then linearly scaled as follows:

Given a protein *x_j_*, its associated average concentration 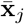 and its measured steady-state distribution *D_x_j|ss__*, a scaling factor *z_j_* was calculated as

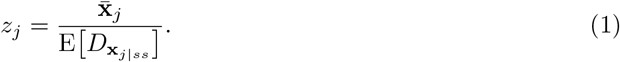

This implementation represents a linear scaling factor of the experimentally measured steady-state distribution of protein *x_j_* such that the mean of the steady-state distribution E|*D_x_j|ss__*] is equal to the population average 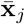 estimated from the quantitative abundance measurements in MaxQB. For all experimental measurements, e.g., after a perturbation, each total protein *j* was scaled using the corresponding *z_j_*.

Steady-state levels of phosphoproteins clearly cannot not be greater than those of the corresponding total protein pools. Additionally, if steady-state phosphoprotein levels are too high relative to total protein, the relative increase in phosphorylation levels in the model would be capped due to a lack of unphosphorylated protein. To avoid these issues, phospho-protein distributions were scaled such that the average steady-state value of a phospho-protein was a fractional value of the total protein level. The additional scaling factors for active MEK, ERK and p90RSK were 0.05, 0.025 and 0.025, respectively. These values were chosen to reduce the number of cells in violation of the “active cannot be greater than total” constraint at steady state; specifically, after scaling, we used the constraint that active protein divided by total protein cannot be greater than 0.3 for MEK, ERK and p90RSK. In the special case that individual cells violated this constraint, these cells were excluded from the sample for analysis (54 of 16,128 cells or 0.34 percent of cells).

**Table S2:** Average expression of total protein 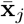 in HEK cells (Boss et al., 2013) used for calculation of mean protein scaling factor *z_j_*.

| Protein | Log10 Expression |
| --- | --- |
| ERK1/2 | 8.213269198 |
| MEK1/2 | 8.344192782 |
| RPS6K1-3 | 7.512659245 |
| GAPDH | 8.673333333 |

The specific implementation of data normalization and scaling may be found in the Matlab file `modellnitializationCellCycleDeath`.

#### Calculation of parameter sensitivities using parameter MMD profiles

To characterize the ability of the data to constrain model parameters, we calculated approximate parameter profiles in a manner analogous to parameter profile likelihoods (Raue et al., 2009). In contrast to classical sensitivity analysis, which calculates the change in model fitness associated with a change in a model parameter value, parameter MMD profiles calculate the change in model fitness associated with a change in a model parameter value that cannot be compensated by any combination of changes to all other model parameters.

Specifically, for any given model parameter *θ_i_* and value 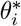, the associated parameter profile point is calculated by fixing 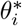 at a value 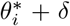, where *δ* ≠ 0, and re-optimizing all other free model parameters *θ_j_,j* ≠ *i*. The full parameter MMD profile for the parameter *θ_i_* is generated by calculating the set of MMD cost values associated with as set of *δ* values *δ* ∈ Δ.

Parameter MMD profiles were calculated for the WT model using 25 linearly spaced samples taken from the parameter-specific ranges described in Supplementary Table S3. Thus, for each parameter *θ_i_*, Δ_*i*_ = `linspace`(LB_*i*_, UB_*i*_, 25). For each profile point 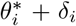, a local search was performed using `fminsearch`, as described above, to determine the minimum associated MMD cost. The “Best Theta” parameter values 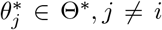 were taken as the initial point for the subsequent re-optimization. Parameter MMD profiles for all model parameters of the WT model are shown in Supplementary Figure S4. Calculation of parameter MMD profiles is implemented in `profileMMD_ALL`.

**Table S3:** Lower (LB) and upper (UB) bounds and associated space of the parameter for calculation of parameter MMD profiles. Profiles were calculated using 25 linearly spaced samples between LB and UB.

| Theta | LB | UB | Space |
| --- | --- | --- | --- |
| $k_f$ | -5 | 5 | $\log_{10}$ |
| $g_D$ | -0.1 | 10 | *Linear* |
| $g_{FB}$ | -0.1 | 10 | *Linear* |
| $k_3$ | -20 | 20 | $\log_{10}$ |
| $g_1$ | -0.1 | 10 | *Linear* |
| $g_2$ | -0.1 | 10 | *Linear* |
| $h_1$ | -0.1 | 10 | *Linear* |
| $k_5$ | -15 | 15 | $\log_{10}$ |
| $g_3$ | -0.1 | 10 | *Linear* |
| $g_4$ | -0.1 | 10 | *Linear* |
| $h_2$ | -0.1 | 10 | *Linear* |
| $k_7$ | -15 | 15 | $\log_{10}$ |
| $g_5$ | -0.1 | 10 | *Linear* |
| $g_6$ | -0.1 | 10 | *Linear* |
| $h_3$ | -0.1 | 10 | *Linear* |
| $I_f$ | 0 | 10 | *Linear* |
| $\tau$ | 0 | 10 | *Linear* |

**Figure S4:**
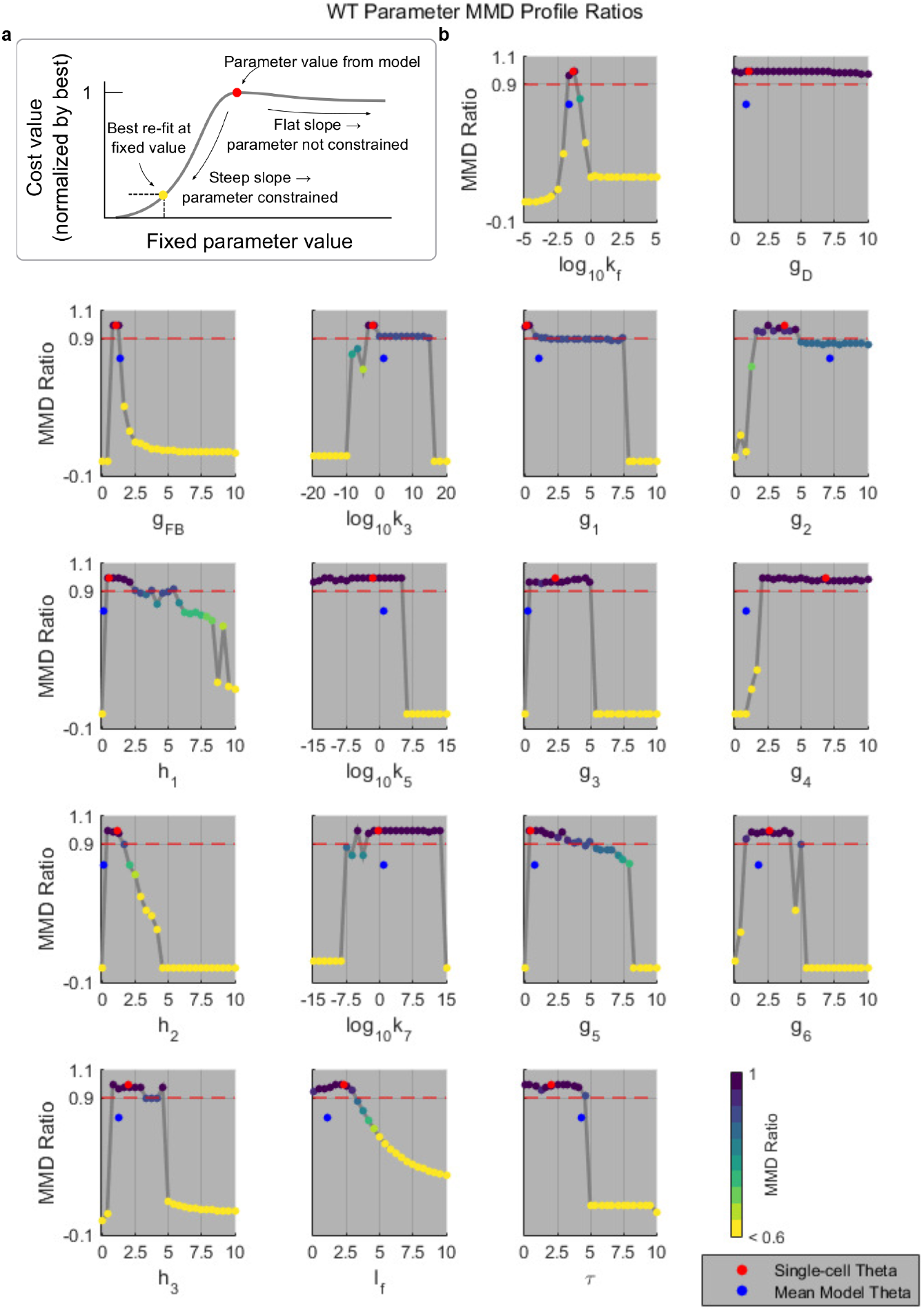
Parameter MMD Profiles. (**a**) Representative illustration of parameter MMD profiles for single-cell model. Each dot represents a point at which the model was re-optimized for the fixed individual parameter value. The color of each point reflects the normalized cost function value. (**b**) Parameter MMD profiles for single-cell model. Red dot: parameter value inferred using DISCO for the WT single-cell model (as in Fig. 3). Blue dot: parameter value inferred using the population-mean modeling approach (as in Supp. Fig. S3)

#### Steady-state-based inference of latent cell-to-cell variation

In real applications of DISCO, we typically do not know the full system structure of 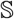 and/or cannot measure all relevant state variables. Expressed differently, the dimensionality of the model **X**\* or the measurements **D**\* is less than the full system. In either case, the corresponding ODE model 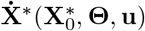 that approximates 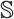 can naturally not account for some source(s) of variation in initial cell states that may influence the system dynamics, and the inference procedure may fail to reproduce the variation in system dynamics characterized by snapshot measurements. If the unknown/unmeasurable model variables change on a time scale qualitatively separable from the dynamics being studied, however, it is possible to infer certain characteristics of the cell-to-cell variation in the unknown or unmeasured variables. This inference is made possible by allowing some model parameters to vary across cells, and by determining the values of these parameters by analytical solution of the steady-state equations using experimental measurements.

Specifically, we take a parameter subset **Φ** ⊆ **Θ** and allow these parameters (**Φ**_*k*_) to vary across each cell *k* in a population of *K* cells, while keeping the remaining parameters the same for all cells. For a given estimation of population parameters **Θ**, the individual cell values of **Φ**_*k*_ are algebraically determined by the steady-state solution of the ODE system using the current estimates of the population parameters **Θ** and steady-state measurements 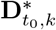 of cell *k* ∈ *K*.

We use an example from our model to illustrate the procedure. We begin with the following differential equation for ERK activation:

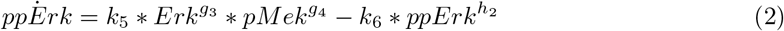

Because the kinetic order parameters *g*_3_, *g*_4_ and *h*_2_ are directly related to physical properties of system components explicitly represented in the model as measured state variables, we assume they do not change across the cell population (thus, {*g*_3_, *g*_4_, *h*_2_} ∈ **Θ**). If our reaction scheme for ERK activation were complete, we would expect that the rate constant parameters *k*_5_ and *k*_6_ to be the same across the population. Our reaction scheme, however, is an approximation of the true reaction structure and does not include all possible contributors to the reaction, such as mono-phosphorylated ERK in the forward reaction and ERK phosphatase(s) in the reverse reaction. For our given experimental conditions, we begin with the assumption that our model explicitly captures the primary state variables that change their state on the same time scale. For instance, we assume that mono-phosphorylated ERK may rapidly equilibrate and the phosphatase is constitutively active. If this assumption is incorrect, our model is unlikely to be able to fit the data and we know our model structure must be updated. By contrast, if our assumption is a reasonablle approximation for the experimental conditions of interest, we may use the parameters *k*_5_ and *k*_6_ to capture the sources of latent cell-to-cell variation. Thus, *k*_5_ and *k*_6_ are included in the single-cell parameters set **Φ**_*k*_. The single-cell parameter values *k*_5*k*_, taken across all cells, capture variation in mono-phosphorylated ERK and any unaccounted ERK activators, while *k*_6,*k*_ captures the variation in ERK phosphatase and any unaccounted deactivators of ERK. Given some guess of **Θ**, which may be set based on biological knowledge or determined by the optimization algorithm, we may use the model structure from equation (2) and our measurement of cell *k* at steady state 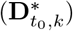 to infer some properties of single-cell parameter values *k*_5*k*_ and *k*_6,*k*_ by solving the steady state equation:

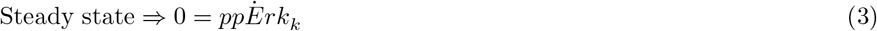

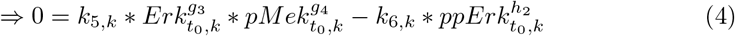

and, by rearranging:

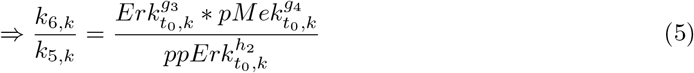

The parameters *k*_5*k*_ and *k*_6,*k*_ are non-identifiable at the steady state due to the model structure, and we only identify their ratio. If one parameter, say *k*_5,*k*_, is fixed, however, the other (*k*_6,*k*_) is fully determined by the model structure, the current population-level parameter values **Θ**, and the constraint that the system must be at a steady state with the current (measured) combination of state variables. The fixed parameter *k*_5,*k*_, while non-identifiable at the steady state, serves to scale the speed of the forward and reverse reactions and can therefore be inferred from the time course measurements of the system dynamics. If we assume the free parameter *k*_6,*k*_ to be responsible for capturing all the cell-to-cell variation in the ERK activation/inactivation reaction, then parameter *k*_5,*k*_ can be added back to the population-level parameter set:

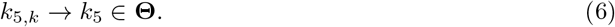

The result is that, first, the number of unknown kinetic parameters for the reaction system describing the change in active ERK is decoupled from the number of cells and, secondly, that the union of single-cell parameter values *k*_6,*k*_ for all cells k in the sample defines a distribution *k*_6_ that captures the unmeasured/unknown sources of cell-to-cell variation in the reaction scheme. Given additional measurement features, we may be able to determine further contributors to the reactions of ERK activation by looking at single-cell correlations between these features and the inferred distribution of *k*_6_ (*k*_6,*k*_ across all *K* cells).

#### Additional supplementary figures

**Figure S5:**
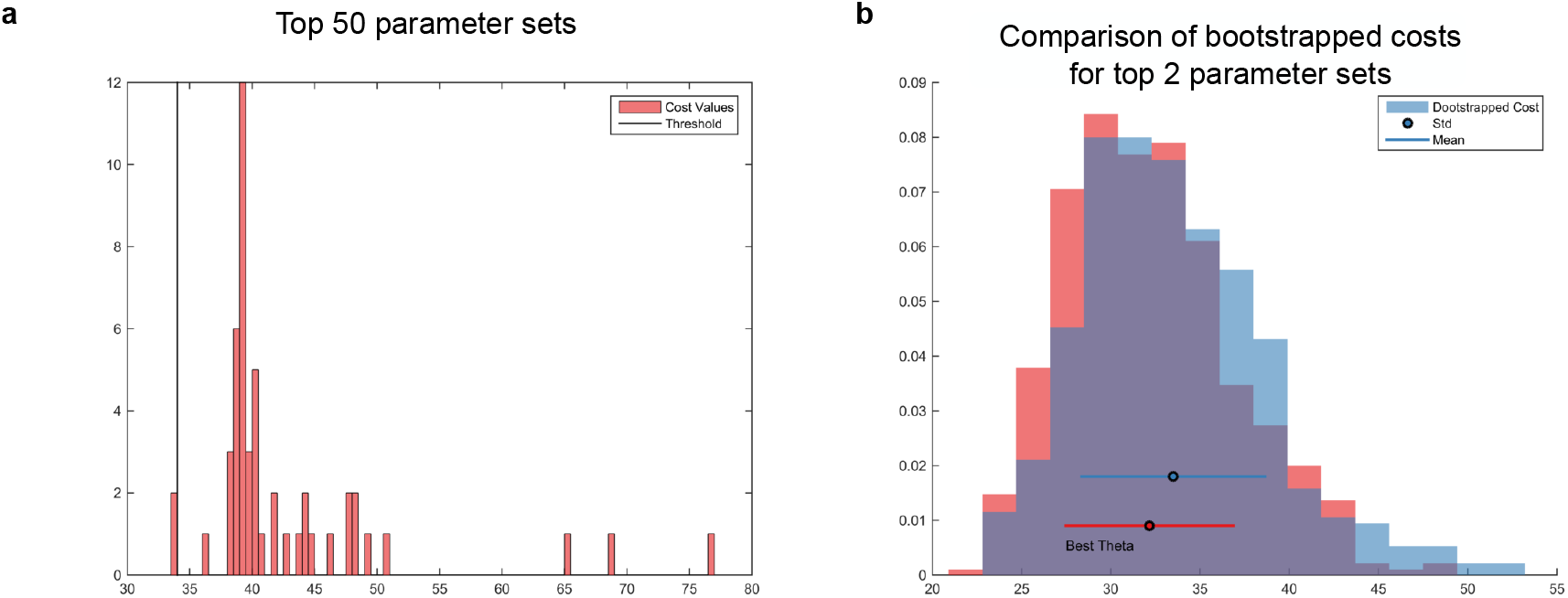
Selection of WT model parameter set. (**a**) Distribution of WT model (MMD) cost for the top 50 (out of 50,000 initial parameter sets) after local search refinement. Parameter sets with model cost less than the threshold were selected for bootstrapped evaluation of robustness. (**b**) Distribution of WT model costs based on repeated sampling (without replacement) and simulation of cell subsets for the parameter sets below the threshold in panel (a). Shown are the mean (dot) +/− standard deviation (line) for each candidate parameter set. The parameter set labeled “Best Theta” was selected for subsequent model analysis.

**Figure S6:**
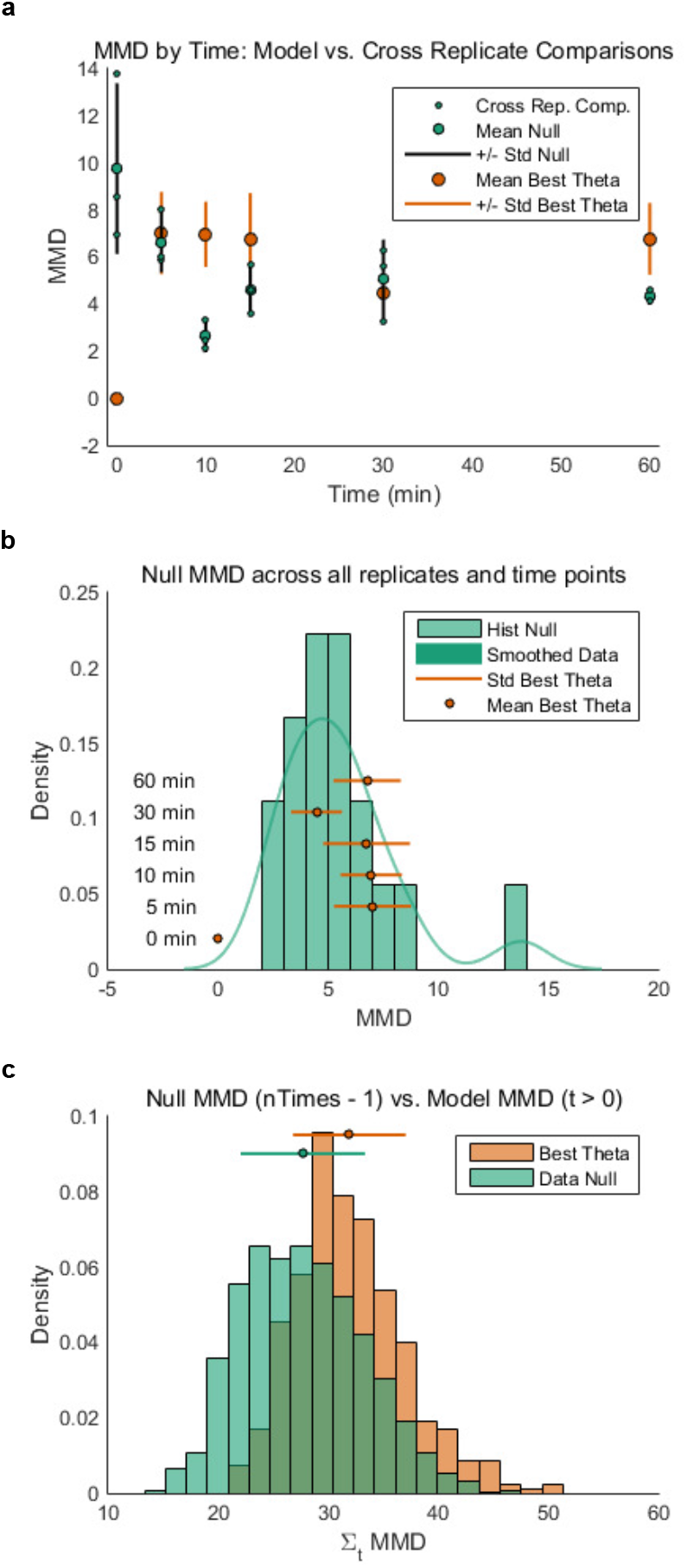
Quality control of bootstrapped WT model. (**a**) Bootstrapped MMD cost of WT model and experimental cross replicate comparisons by time point. (**b**) Bootstrapped MMD cost of WT model and experimental cross-replicate comparisons across by time point. Standard deviation line is mean +/− standard deviation. (**c**) Bootstrapped MMD cost of WT model and experimental cross-replicate comparisons summed over all time points (model objective function as in Figure 3b).

**Figure S7:**
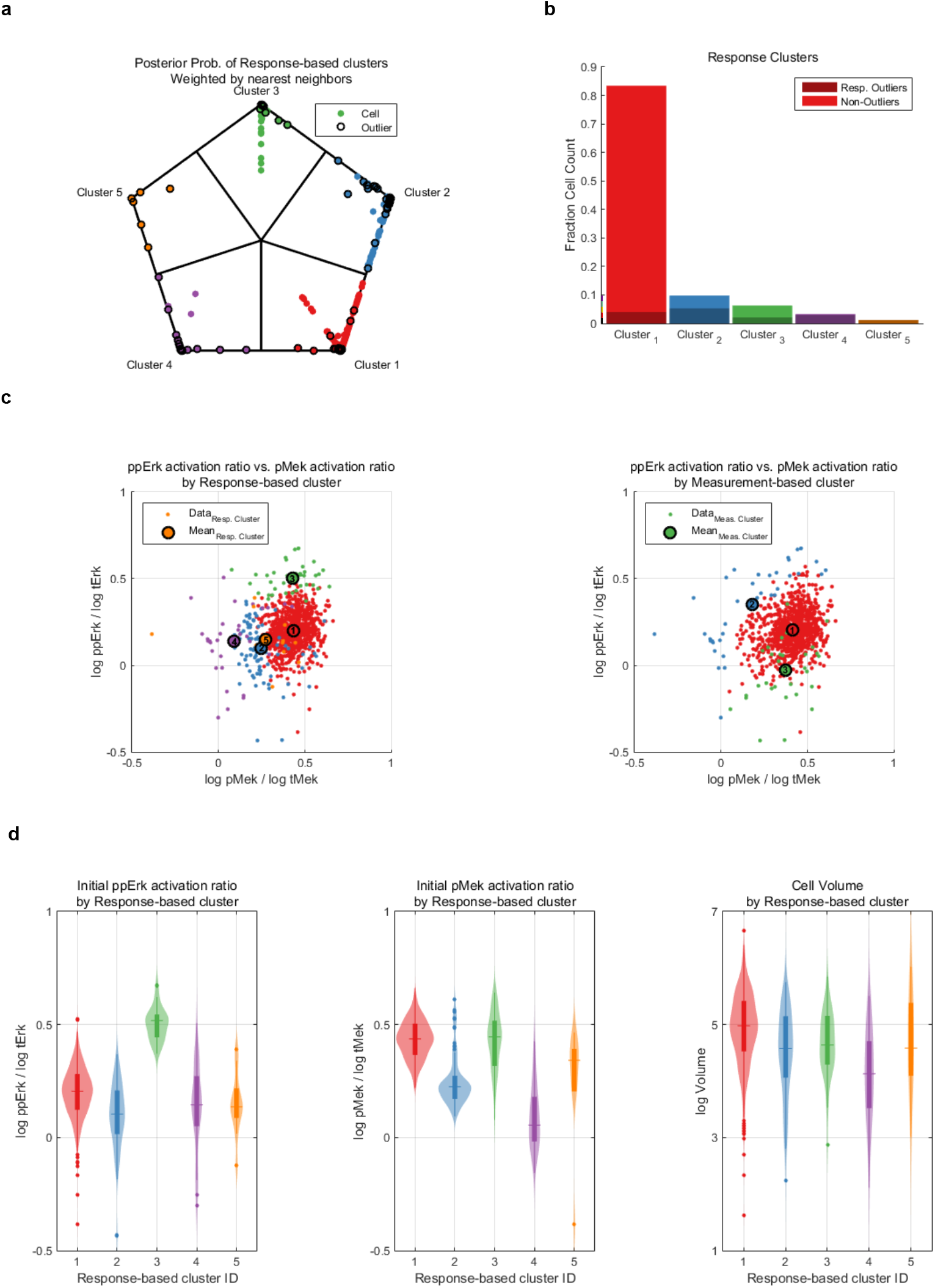
Subpopulation analysis. (**a**) Weighted posterior probability of cell assignment to top 2 nearest neighbor response clusters. Response-based outlier cells are marked with black circles. (**b**) Distribution of cells and response based outliers by cluster ID. (**c**) Initial activation ratio of MEK (pMEK/total MEK) versus ERK (ppERK/total ERK) with cells colored by response-based (left) or measurement-based (right) cluster IDs. Large circles indicate projected cluster centers. (**d**) Violin plots of response-based clusters by initial activation ratio of MEK, initial activation ratio of ERK and cell size.

**Figure S8:**
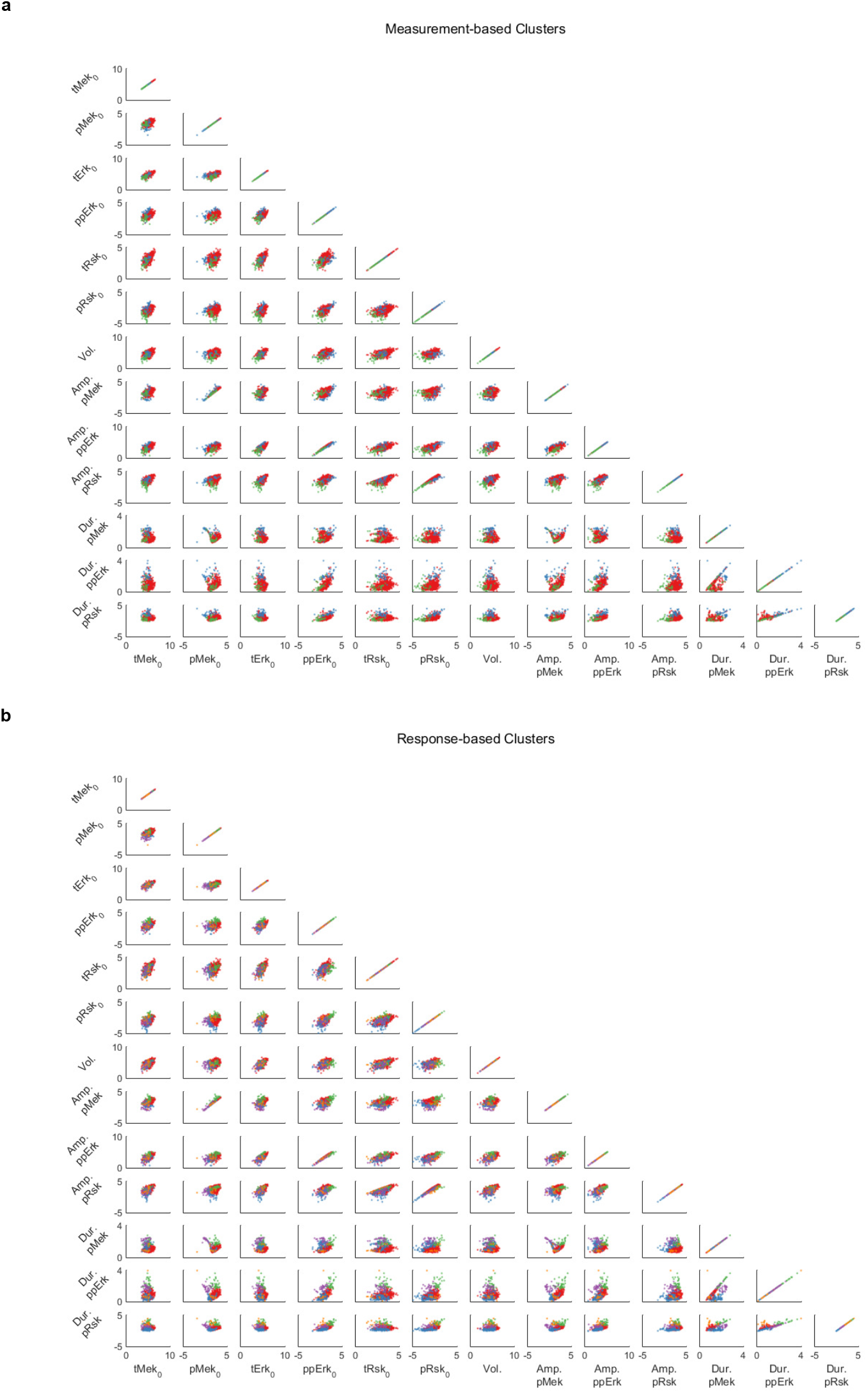
Measurement-based and response-based clusters on all cell measurement and response features. (**a**) Scatter plots of all cell response and measurement features colored by measurement-based cluster ID. (**b**) Scatter plots of all cell response and measurement features colored by response-based cluster ID.

**Figure S9:**
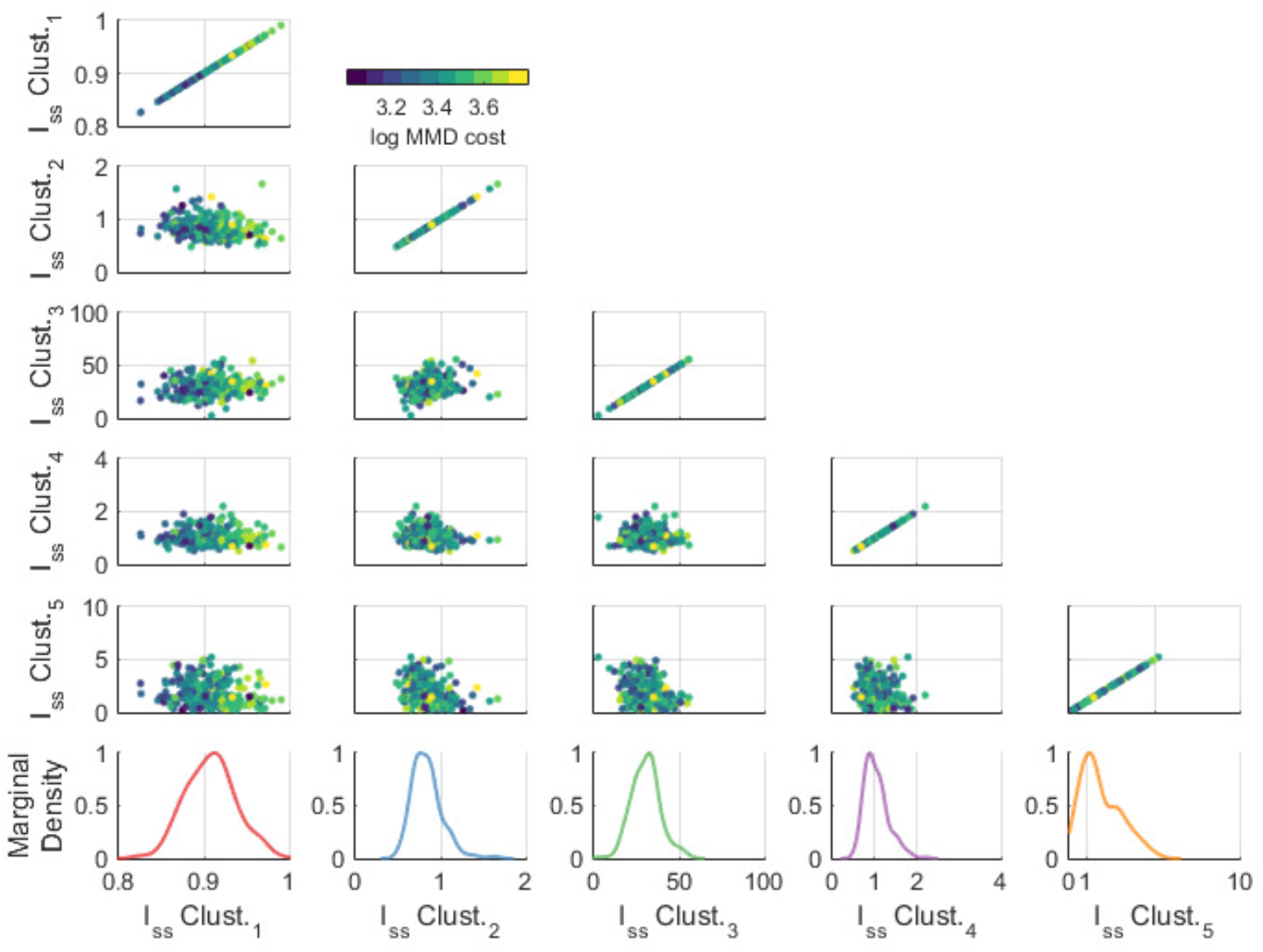
Steady-state input *(Iss)* distribution by response-based cluster. Each point corresponds to local optimization of response-cluster dependent *Iss* for an independent subsample of cells (sampled without replacement from data). Points are colored by corresponding log of model (MMD) cost. Marginal distributions of *Iss* are colored by response-cluster ID.

**Figure S10:**
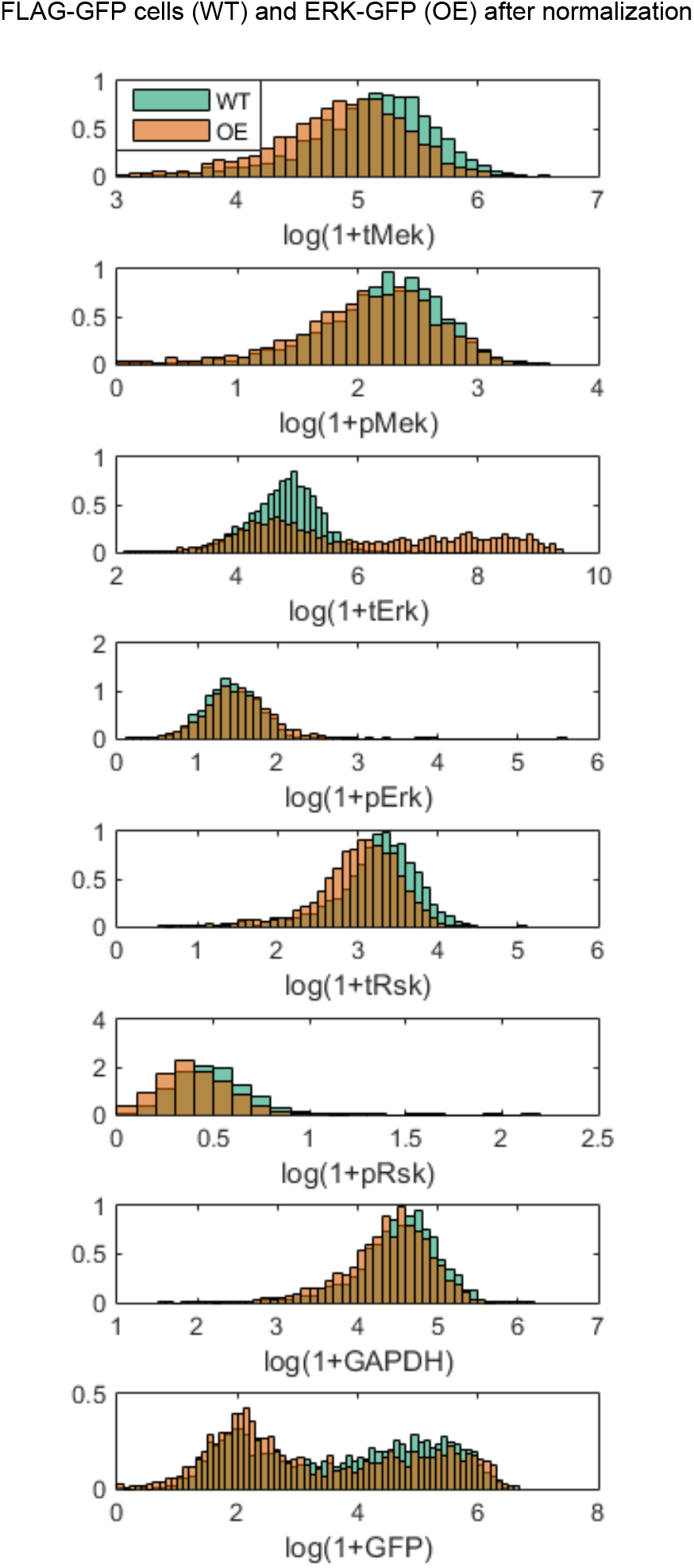
Marginal distributions of ERK2-GFP overexpression condition at steady state after normalization to FLAG-GFP (WT) measurements.

**Figure S11:**
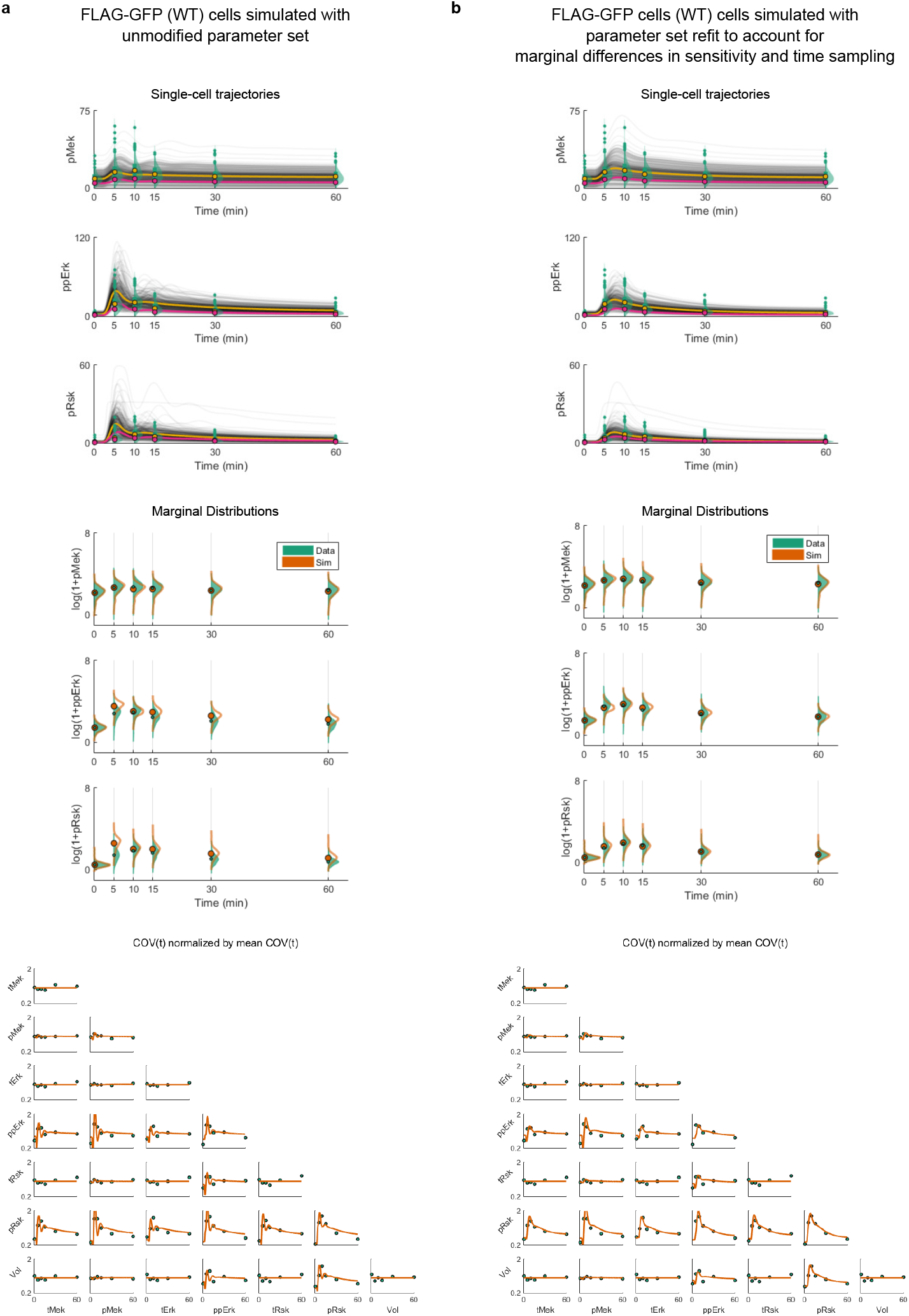
Example of model simulations of FLAG-GFP data before and after adjusting kinetic parameters to account for machine sensitivity and marginal differences in experimental timing. (**a**) Single-cell simulations (top), marginal distributions (middle) and normalized covariance over time (bottom) of simulations using FLAG-GFP data before adjusting parameters from WT model as in Figure 3 (**b**) Single-cell simulations (top), marginal distributions (middle) and normalized covariance over time (bottom) of simulations using FLAG-GFP data after local re-optimization of model parameters. For single-cell simulations: Yellow circles and line are the mean of the data and simulations, respectively; note linear y-axis. For marginal distributions, the data are green and single-cell model simulations orange; note log y-axes. For normalized covariance:, the covariance between model components over time for measurements is given as green circles (by replicate) and model simulations (orange line). In each plot, covariance has been normalized by the average covariance over time.

**Figure S12:**
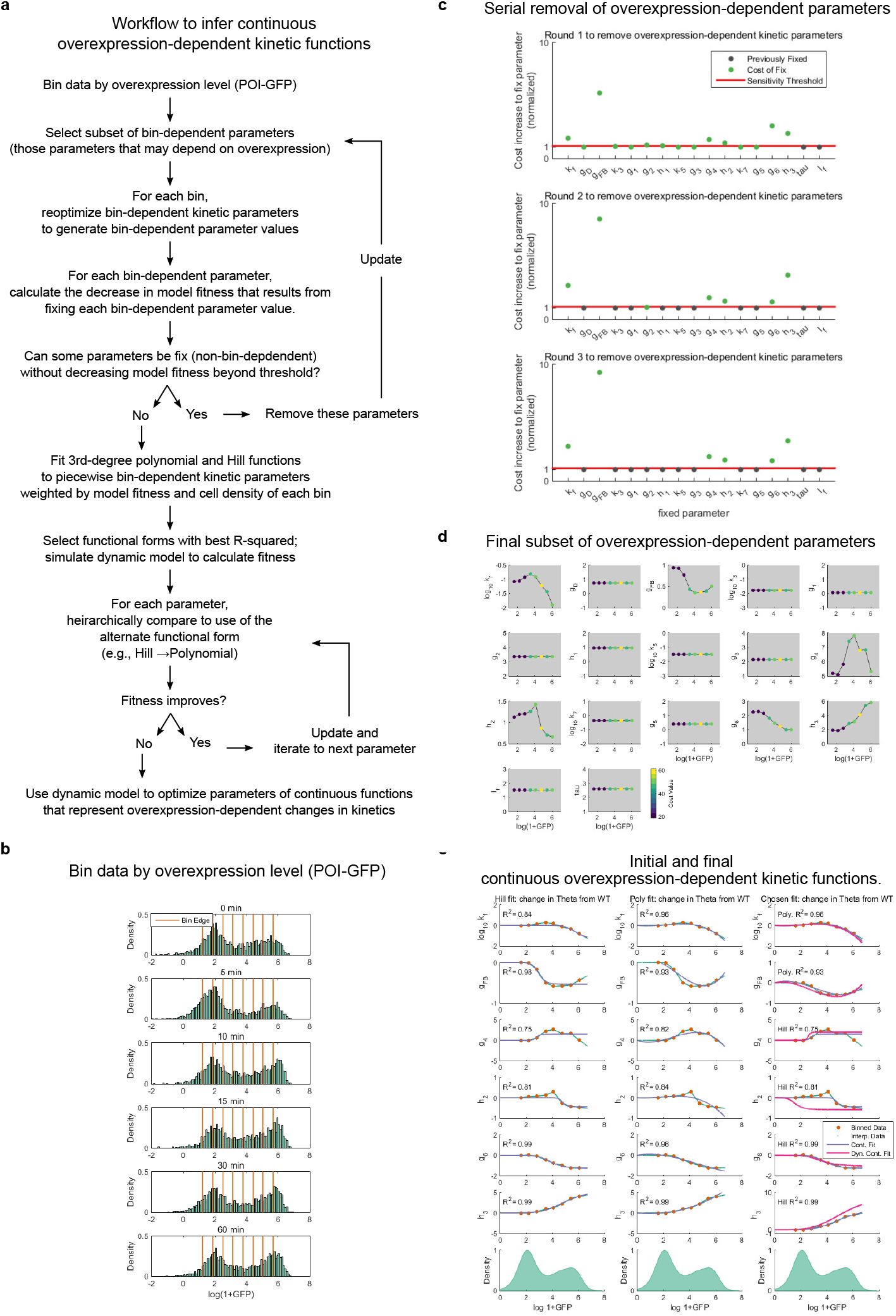
Overview of algorithm for overexpression-dependent kinetic parameter inference and intermediate results. (**a**) Graphical description of the DISCO-based algorithm to infer continuous expression-dependent parameter functions. (**b**) Binning of ERK2-GFP by time point. (**c**) Serial reduction in expression-dependent model parameters (green dots) by algorithmic step. Threshold (red line) is calculated as described in *Methods.* Parameters that are fixed based on the previous step are represented by gray dots. (**d**) Piecewise-linear functional representation of model parameter values by bin. Single-cell model (MMD) cost, as determined by application of DISCO to each bin, is represented by the dot color. (**d**) Illustration of initial fit (violet line) of a Hill function (left) and polynomial (middle) to the piecewise-linear expression-dependent parameter values (orange dots) from (c), interpolated at the single-cell level (green dots) as described in Methods. The final functional form chosen to represent expression-dependent parameter values with parameters optimized using DISCO (magenta line).

**Figure S13:**
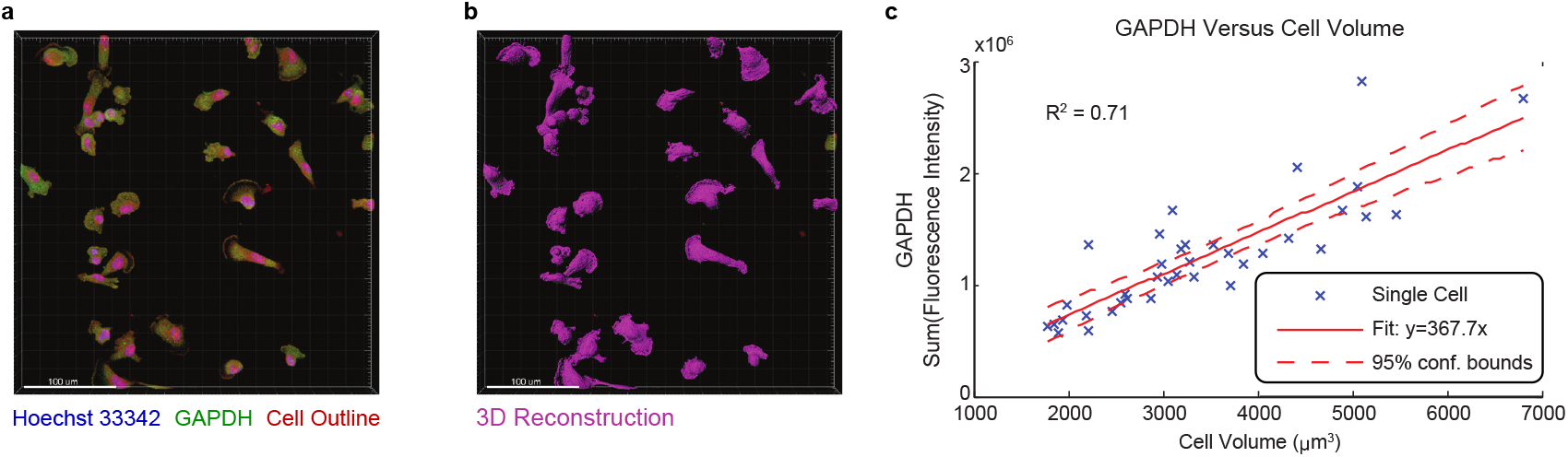
Correlation between GAPDH and cell volume. (a) Confocal images of cells stained for GAPDH (green). Nuclear staining by Hoechst 33342 (blue) and Alexa Fluor 647 carboxylic acid succinimidyl ester to determine cell outline (red). (b) Three-dimensional reconstruction of cells based on cell outline staining. (c) Regression of GAPDH versus reconstructed cell volume.

**Figure S14:**
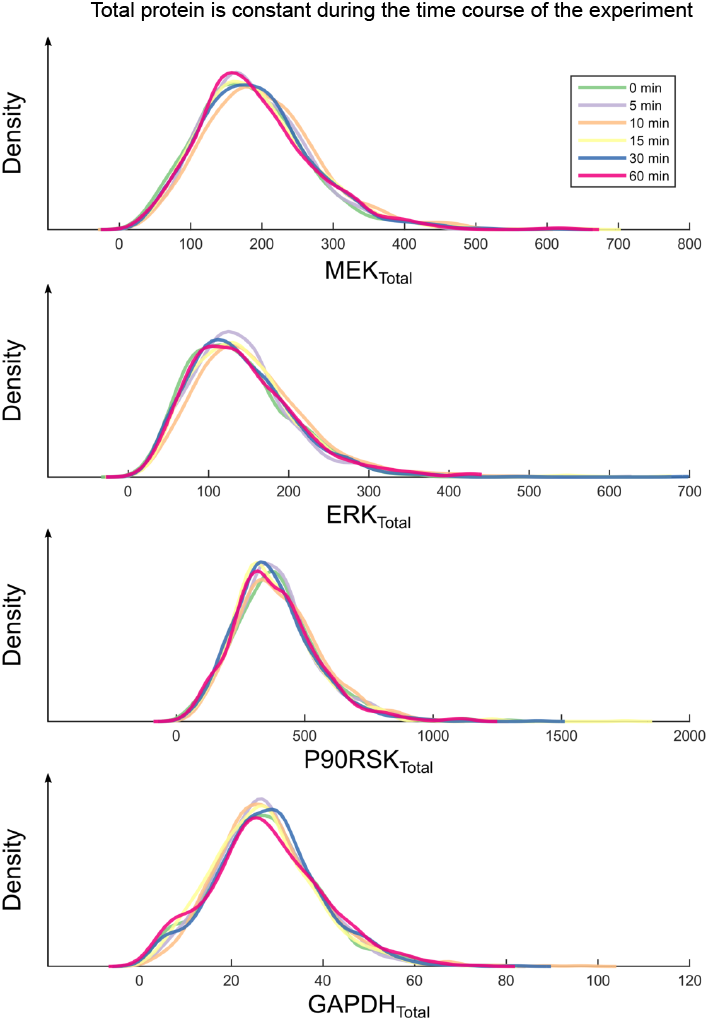
Distributions of total proteins remain constant across time points sampled during the experiments.

**Table S4:**
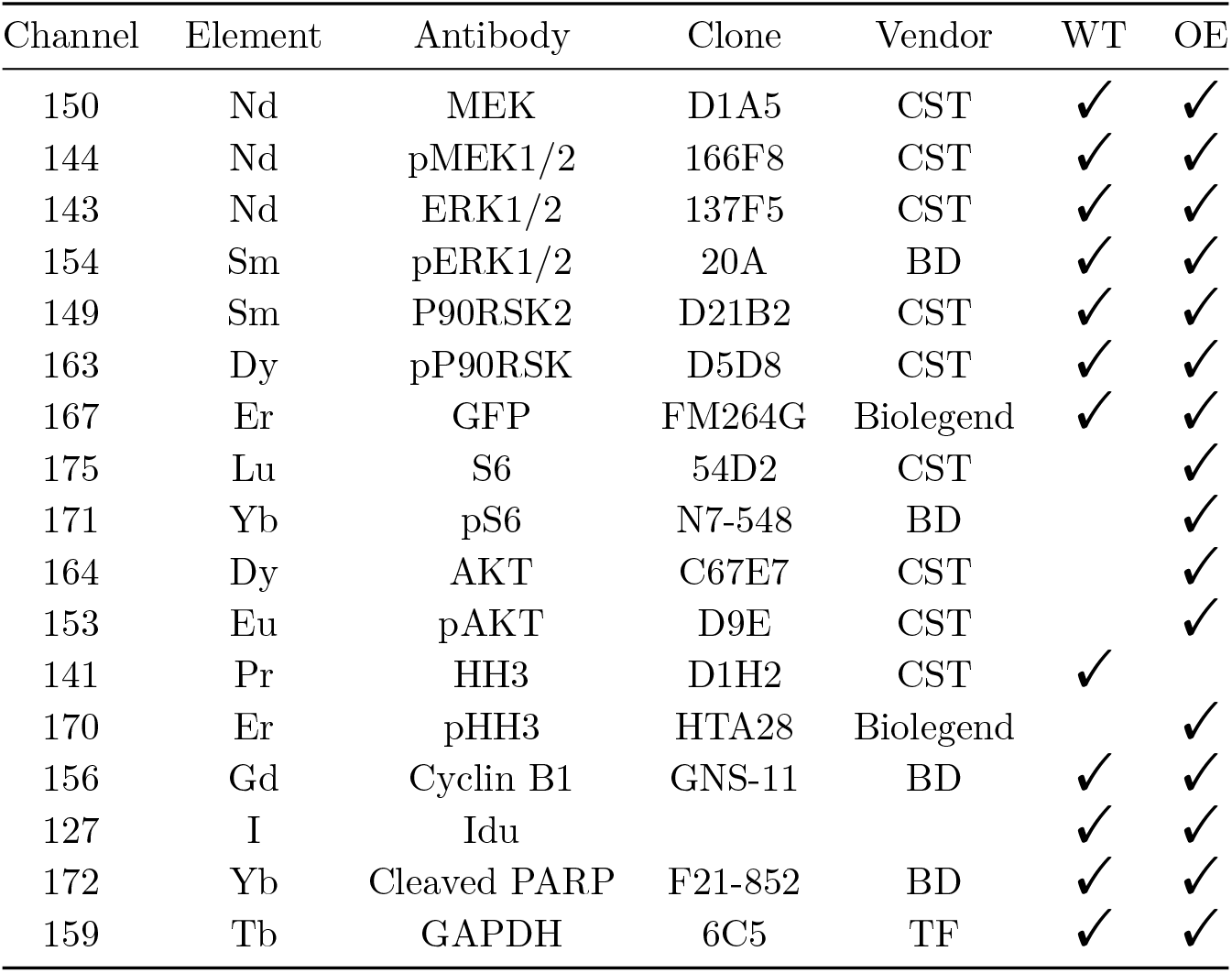
Antibody panel for wild type (WT) and overexpression (OE) experiments. TF: Thermo Fisher; CST: Cell Signaling Technology; BD: BD Biosciences.

